# Global-scale quantification of responses to anthropogenic stressors in six riverine organism groups

**DOI:** 10.1101/2024.07.06.602319

**Authors:** Willem Kaijser, Christian Schürings, Andrea Schneider, Sebastian Prati, Michelle Musiol, Franziska Wenskus, Verena S. Brauer, Lena Feldhaus, Cornelia S. Wagner, Rike Bayer, Iris Madge Pimentel, Sebastian Birk, Brauns Mario, Louisa Dunne, Julian Enss, Luan Farias, Christian K. Feld, Svenja M. Gillmann, Kamil Hupalo, Stephen E. Osakpolor, Sarah Olberg, Christian Schlautmann, Jessica Schwelm, Nicole E. Wells, Bernd Sures, Ralf B. Schäfer, Daniel Hering

## Abstract

Rivers globally are impacted by numerous anthropogenic stressors, including water pollution, habitat degradation, and climate change, which collectively stress biodiversity and ecosystem functioning. This study systematically reviews and analyses published and unpublished data to understand how five aquatic organism groups (bacteria, algae, macrophytes, invertebrates, fish) respond to seven common stressors (salinization, oxygen depletion, fine sediment enrichment, temperature increase, flow modifications and nitrogen or phosphorus enrichment). Using an analytical framework that includes Generalized Linear Models (GLMs) and Robust Bayesian Meta Analysis (RoBMA), we extracted data from 143 relevant datasets out of 29,749 screened articles. Our results reveal a negative relationship between invertebrates and salinity, fine sediment enrichment, and temperature increase, while fish respond positively to increased oxygen levels and temperature. Bacteria and algae show variable responses, with algae positively associated with nitrogen. The findings highlight strong variability in stressor-response relations across organism groups and stressor types, and emphasize the need for more targeted studies on underrepresented groups like macrophytes and microorganisms. This analysis enhances the predictive understanding of stressor impacts on riverine biodiversity, informing future river ecosystem management and restoration efforts.

## 1. Introduction

Worldwide, most rivers are affected by a multitude of anthropogenic stressors, including human-induced physico-chemical water pollution, hydro-morphological habitat degradation, and impacts associated with climate change, such as hydrological and temperature modifications (Dudgeon et al., 2006; Fouchy et al., 2019; Grill et al., 2019; Reid et al., 2019; Ríos-Touma and Ramírez, 2019; Vörösmarty et al., 2010; Woodward et al., 2010). Multiple stressors are therefore a pervasive phenomenon that affects biodiversity and the functioning of river ecosystems. In many cases, both natural disturbances (such as droughts) and anthropogenic stressors (such as eutrophication) do not cause direct biological effects, but rather alter environmental variables such as temperature, oxygen concentration, substrate composition and flow, all of which affect biota both individually and in combination, thus determining the occurrence and survival of organisms. Consequently, anthropogenic stressors typically reduce biodiversity and negatively impact ecosystem functioning (Peipoch et al., 2015; Reid et al., 2019).

Over the past two decades, numerous case studies have explored the impacts of multiple stressors on aquatic organisms and ecosystems. More recently, efforts have intensified to categorize and classify these studies by various criteria, such as ecosystem type (Nõges et al., 2016), organism group (Jackson et al., 2016), reversibility (Sabater et al., 2019), stressor duration (Perujo et al., 2021), and temporal overlaps of stressors (Jackson et al., 2021). These attempts, however, did not necessarily lead to a deeper understanding of the mechanisms that guide multiple stressor response relations. Only recently, some concepts emerged, capable of classifying the mechanisms of stressor interactions (Jackson et al., 2021; Pirotta et al., 2022), partly associated with practical management guidance (Spears et al., 2021; Vos et al., 2023). However, we are still far from reliably predicting the biotic response based on multiple stressor increase or release.

The basis for the analysis of *multiple* stressors-response relations is a quantified understanding of the magnitude of change between *individual* stressors-response relations, and even this has not yet been achieved. While many individual case studies explored stressors-response relations such as flow alteration, salinization and eutrophication on individual species, communities and ecosystems, a generalization of their impacts across geographical regions and river types has to our knowledge not been attempted. The stressors affecting riverine organisms can roughly be grouped into physico-chemical and hydro-morphological impairments. Salinity, for instance, is an important stressor affecting organisms’ osmoregulation, however, its study in freshwater systems remains limited (Cañedo-Argüelles et al., 2013). Oxygen depletion has been more extensively explored, revealing negative correlations with invertebrates and fish (Frakes et al., 2021; Jonsson and Jonsson, 2009; Nikinmaa and Rees, 2005). Temperature is an emerging stressor aggravated by climate change, causing both physiological stress and indirectly affecting oxygen concentrations and thereby invertebrates and fish (Frakes et al., 2021; Jonsson and Jonsson, 2009; Reid et al., 2019). Substrate type and flow velocity strongly shape riverine communities via habitat provision, and their modification leads to a general biodiversity decline (Biggs, 1996; Death and Winterbourn, 1995; Hoyle et al., 2017), while nutrients particularly influence bacteria, benthic and periphytic algae (Hering et al., 2006; Kinsman-Costello et al., 2023; Zhao et al., 2022).

The outcome of each individual stressor-response relation is mediated by the coverage of the observed stressor gradients, co-occurring stressors, and the chosen response variables, i.e., biodiversity metrics. These dependencies limit our ability to compare the response of organism groups to stressor gradients. In addition, there is a scarcity of comparative analyses investigating stressor effects on several organism groups simultaneously. We are aware of just four studies that assessed the association of degradation gradients with four to five aquatic taxonomic groups (Hering et al., 2006; Johnson et al., 2006; Johnson and Hering, 2009; Kaijser et al., 2024) and a few more that addressed responses of different taxonomic groups to restoration, including floodplain-inhabiting terrestrial groups (e.g., Jähnig et al. (2009)).

As a consequence of all these knowledge gaps, our forecasting ability remains limited. Despite decades of research, it is not even possible to reliably predict the biodiversity effects of a defined increase in salinity or nutrient concentration in a generalizable way, or to quantitatively estimate which organism group would be affected most strongly. For future endeavors that will target the mechanisms of multiple stressor-response relations, and even for projects restoring river ecosystems, this scarcity of underlying data is an obstacle: if the amount of information or the magnitude of the stressor-response relation is limited or marginal or the uncertainty too large, there is little incentive to mitigate the stressors.

Here we systematically collect and comparatively analyze published and unpublished data on the relationship between the five most commonly explored aquatic organism groups to the intensity of seven pertaining stressors, resulting in the first-ever comprehensive appraisal of riverine organism response to stress. The main questions guiding our analyses are:

1. How are individual organism groups related to the stressor gradients?
2. How does stressor gradient length affect the stressor-response relation?

## 2. Methods

Our study is based on data extracted from a systematic literature search, complemented with additional datasets. All data was subsequently analyzed with Generalized Linear Models (GLMs), the results of which were combined with Robust Bayesian Meta Analysis (Hinne et al., 2020; Hoeting et al., 1999; Maier et al., 2023). We targeted five organism groups: bacteria, algae (including benthic and planktonic algae), macrophytes, benthic invertebrates and fish; and seven stressor types: salinity, oxygen depletion, fine sediment enrichment, thermal modification, flow modification, and nitrogen and phosphorus enrichment. The workflow is illustrated in Fig. 1.

**Figure 1:**
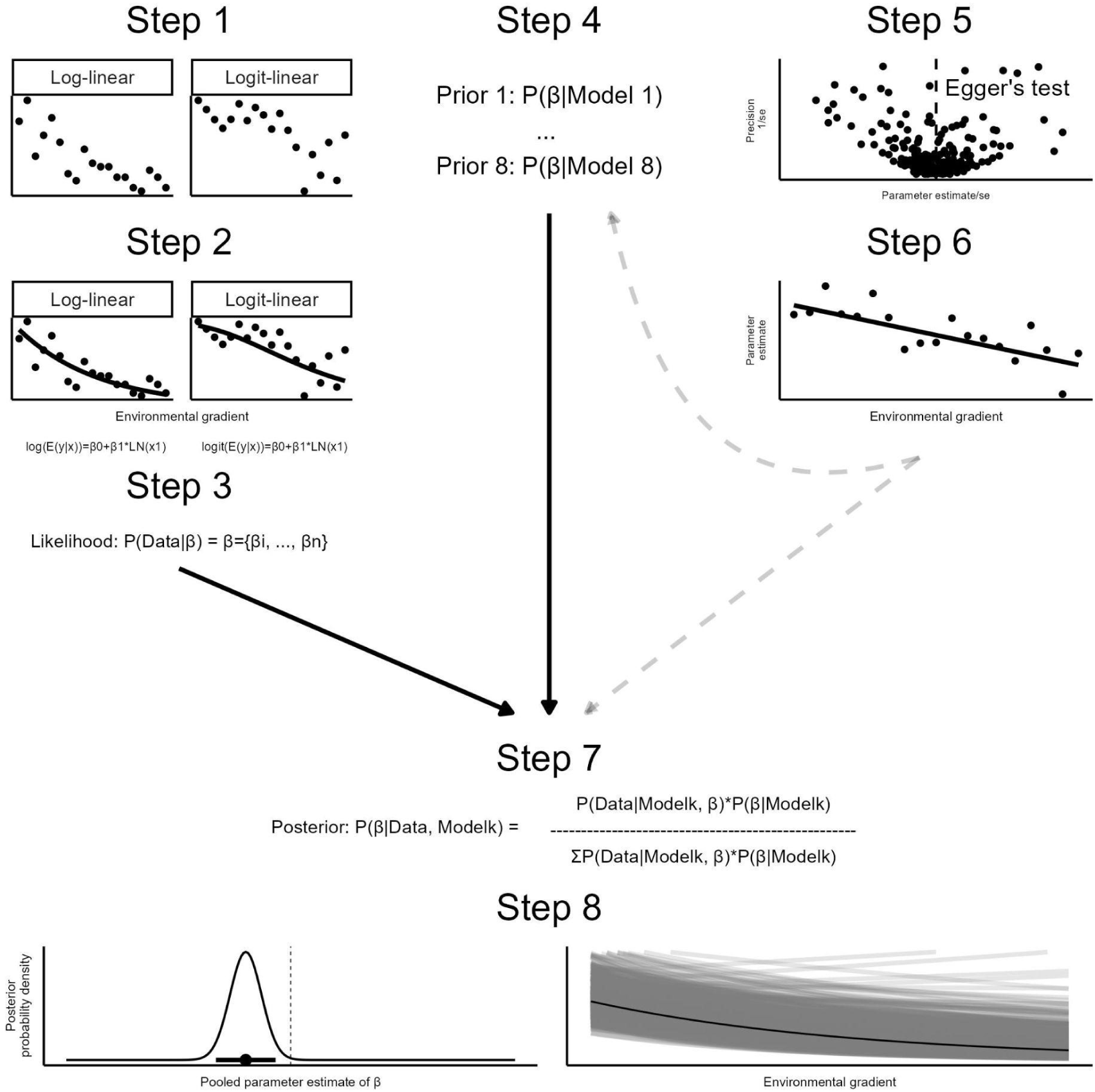
Analytical workflow.

In short, we performed (1) a systematic search in Web-of-Science targeting the five biotic groups in relation to the seven stressors. This yielded 29,749 hits. The response types of interest included taxon richness and evenness (and other fractional responses). We additionally searched non-systematically for datasets in Dryad, GitHub and Zotero (Text S1, Fig. S1). The relevant information was extracted from figures, tables and datasets referred to in the data availability statements. (2) For each extracted dataset, we fitted a GLM using log- or logit-linear models with natural logarithmic transformations of all stressor gradients provided; i.e., for each model, we used up to seven stressor gradients. In log-linear models, the estimated regression coefficient results in the elasticity coefficient, indicating the percentage increase in the response per 1% in the stressor (Woolridge, 2001). For example, an elasticity coefficient of 0.2 indicates a 0.2% increase in response with a 1% increase in the stressor gradient. In logit-linear models, this interpretation loses precision as the deviation from 0 increases. (3) By repeating these steps, we compiled the regression coefficients of each model for a meta-analysis. From here on, the regression coefficients are referred to as ‘parameter estimates’. (4) Eight priors were formulated for each parameter estimate to be used in Robust Bayesian Meta Analysis (RoBMA). (5) The accumulated estimates were checked for bias via Egger’s test. (6) We regressed the parameter estimates resulting from (2) on the stressor gradients, using a Linear Mixed Model (LMM), and the stressor gradient was expressed as the natural logarithm of the coefficient of variation 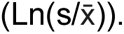 This step answers question 2 (the dependency of the stressor-response relation on the stressor gradient). (7) We performed RoBMA with the information of step 3 (accumulation of the estimates), step 4 (all priors) and step 6 (coefficients of variation; gray dashed lines in Fig. 1). This results in a single pooled parameter estimate for each stressors-response relation per organism group. Finally (8), we displayed the single pooled parameter estimate in posterior density plots and six strong stressor-response relations as simulated lines between stressor and response (Kale et al., 2019). This answers question 1 (how are individual organism groups related to the stressor gradients?). For a full overview of rationale and method, see Text S1.

All models and figures were produced in R (R Core Team, 2023). Models in (2) were fitted using the glmmTMB package version 1.1.8 (Brooks et al., 2017) and GAMLSS version 5.4-20 (Rigby and Stasinopoulos, 2005), and other analyses were performed using JAGS version 4.3.1 (Plummer, 2003) via R2Jags vs. 0.7-1 (Su and Yajima, 2021). All figures were plotted using the packages bezier version 1.1.2, ggplot2 version 3.4.4 and cowplot version 1.1.3 packages (Olsen, 2018; Wickham, 2009; Wilke, 2019).

## 3. Results

Of the 29,749 screened articles and additional retrieved data, 143 datasets were extracted (Fig. S1). Each specific discrete or proportional response of one of the targeted organism groups was regressed on at least one of the stressors, from which the parameter estimate is used in the meta-analysis. Egger’s test (Fig. S2) revealed no bias, except that data extracted from figures predominantly present ‘positive’ results (|Z|>1.96)

Prior to the meta-analysis, all estimates were plotted as box-plots (Fig. 2). The estimates of logit-linear models were slightly larger than for log-linear models. The strongest patterns were observed for the combinations of fish and invertebrates with salinity, fine sediment and temperature. Closer examination of the box-plots for flow reveals that the median for the log-linear models is positive, but negative for the logit-linear models.

**Figure 2:**
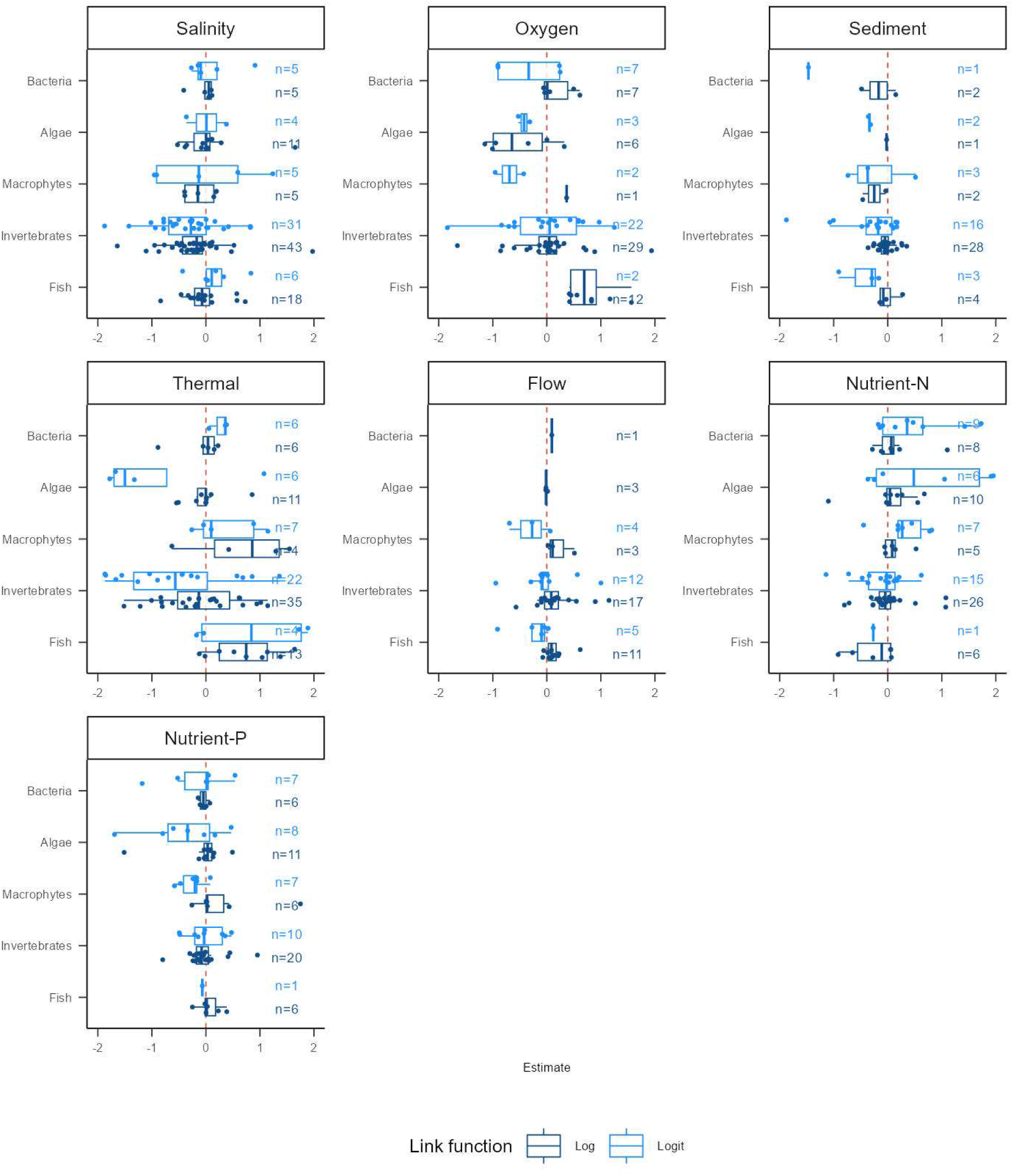
Point estimate (without standard error) for the combinations of organism groups and stressors. The colors of the box plots highlight the estimated parameters from the log- and log-linear models. The box-plot for the logit-model (e.g., oxygen depletion vs. fish) is outside the margins and thus not displayed.

Figs. 3 and 4 present the separate posterior estimates for the log-linear and logit-linear models, respectively. Most organism groups, especially invertebrates and algae, show a negative relationship to salinity in both models. Fish and bacteria display a positive log-linear relation to oxygen, while the relation for invertebrates is less clear. All these patterns are less evident in logit-linear models. Invertebrates are negatively related to fine sediment (%) and temperature in log-linear models, but not in logit-linear ones. For flow modification, all groups, except bacteria and algae, show a positive relation in log-linear models and a clear negative relation in logit-linear models. Algae show a positive relation to nitrogen in both models.

**Figure 3:**
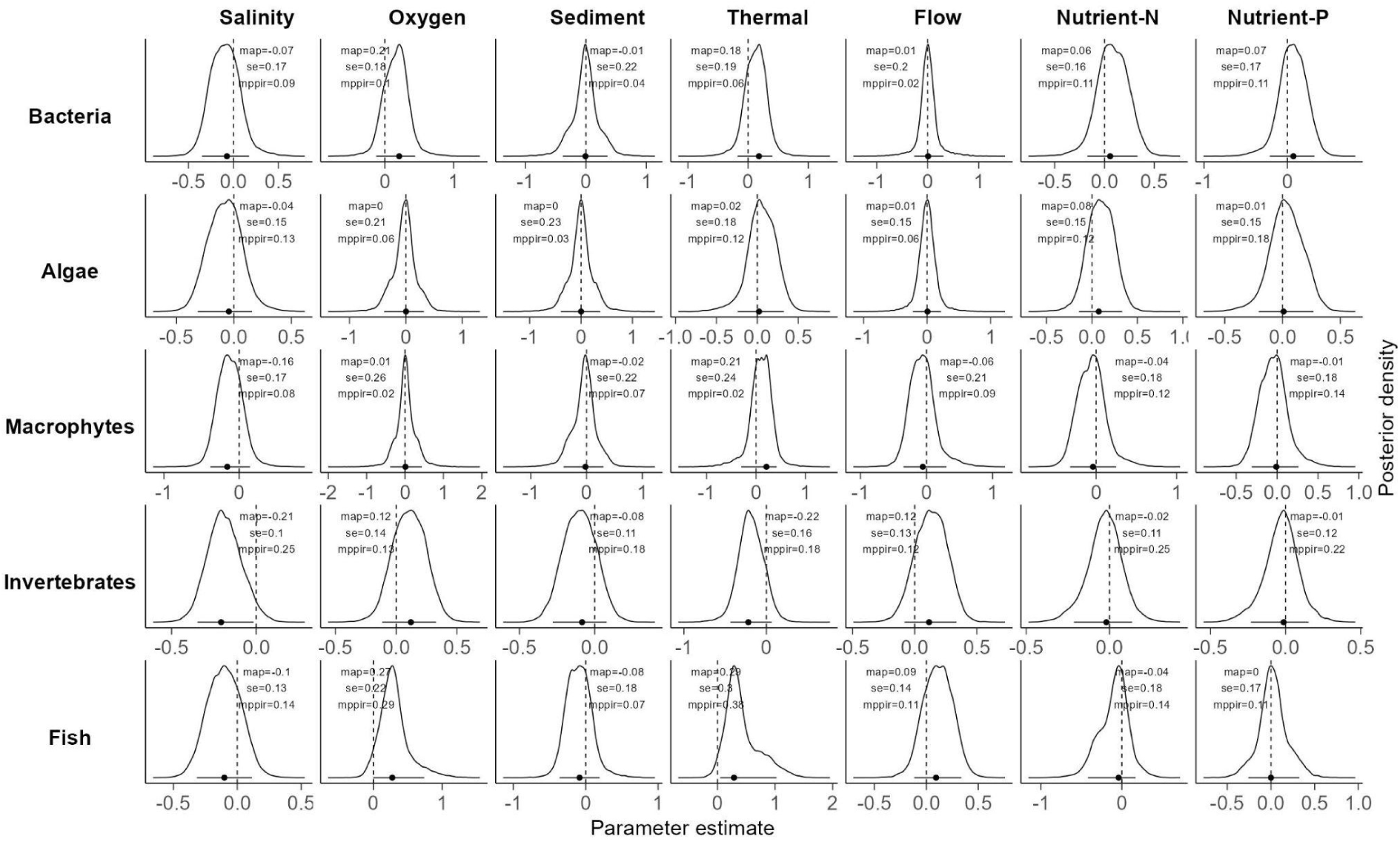
Log-linear models for the response of organism groups (countable responses, such as species number) to stressor gradients. The lower ‘point’ indicates the Maximum A Posteriori (MAP) estimation, indicating the highest density of the posterior estimate, while the error bars indicate the credibility interval at 90%. MAP and SE are given in the figures. Additionally, a metric is given that measures the extent to which the data affects the shift of the prior odds to the posterior odds (mppir). In simple terms, the mmpir indicates how much has been ‘learned’ about the prior guesses from the model, rated as a number between 0 and 1. The value 0 indicates all information the data provided is equal to our prior assumptions, and a value closer to 1 indicates that all information the data provides is shifted to particular prior models.

**Figure 4:**
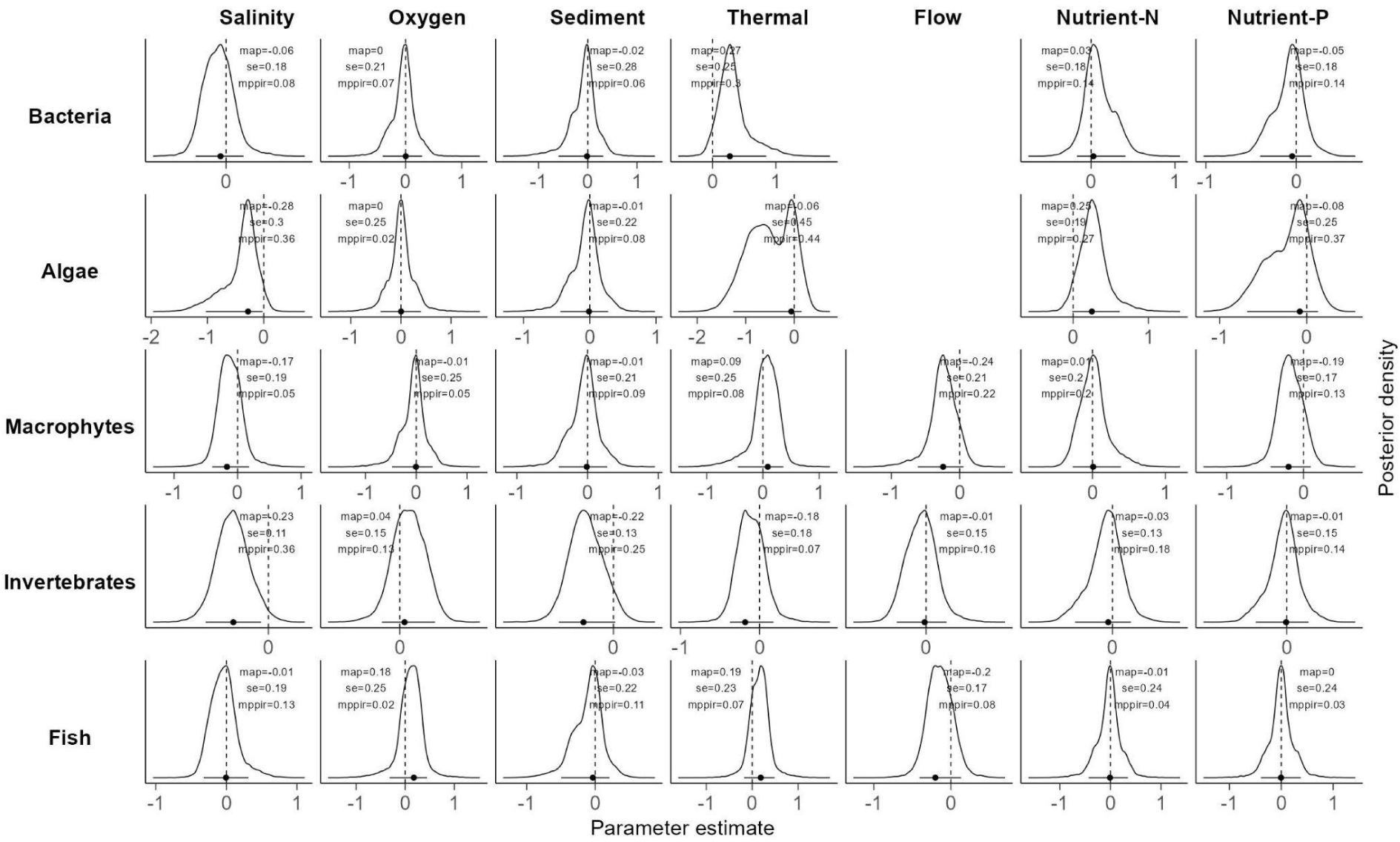
Log linear models for the response of organism groups (relative response variables, such as evenness) to stressor gradients. Compare Fig. 3.

**Figure 5:**
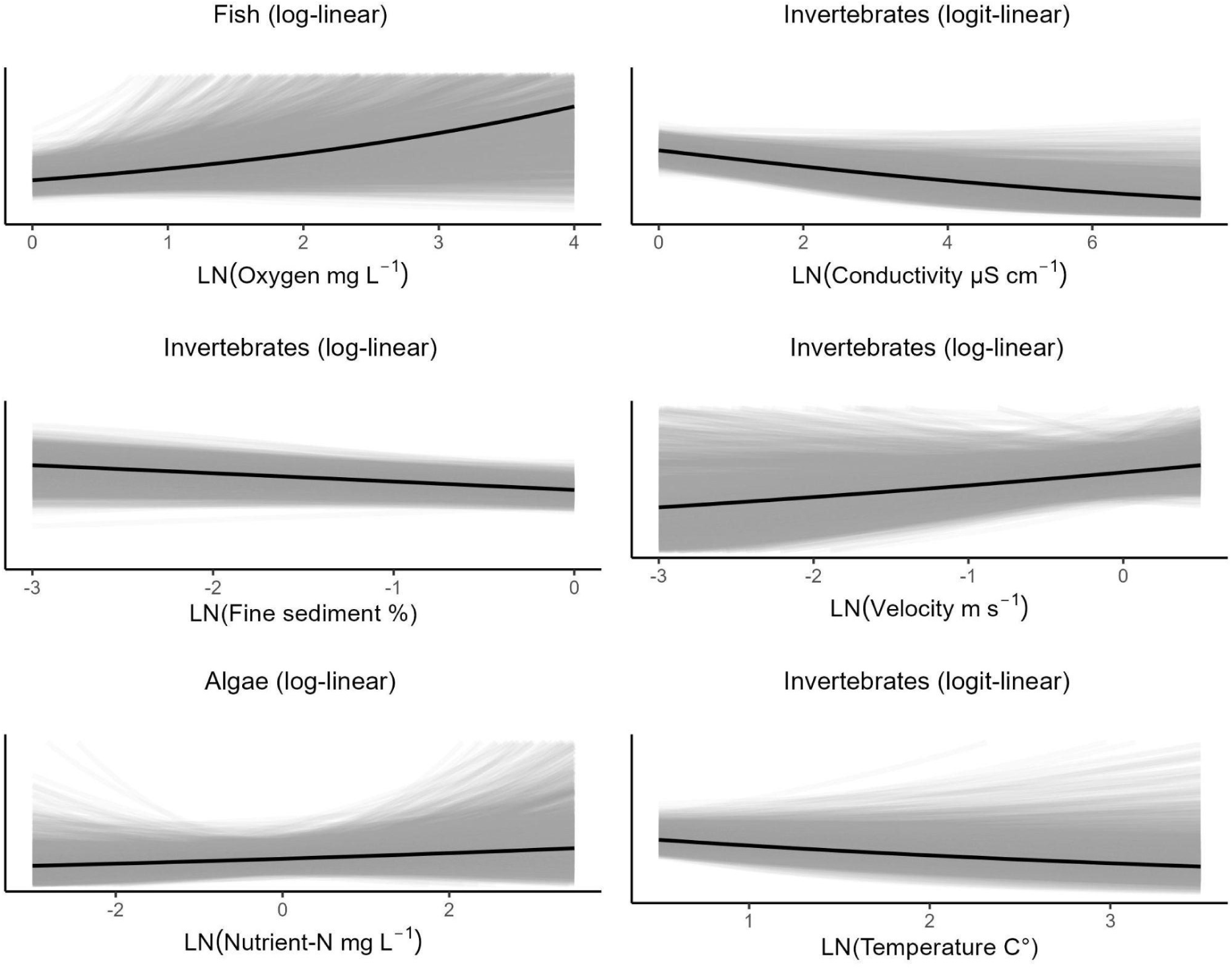
Hypothetical Outcome Plots (HOP) expressing the response of an organism group as a function of the stressor gradient. The y-axis is left unnamed, as responses on different taxonomic levels are included (species, genus, family or order). The dark black line indicates the MAP and the gray lines 1500 simulations from the posterior distribution.

For six stressor-response combinations, Hypothetical-Outcome-Plots (HOPs) were generated, visualizing the expected outcomes of Fig. 3 and 4 given a stressor increase. Fish respond positively to oxygen increase, invertebrates to flow, and algae to Nutrient-N. Strong negative patterns are emerging for the response of invertebrates to conductivity, fine sediment and temperature.

The estimated parameters decreased with an increase in the coefficient of variation (Fig. 6), providing support for the alternative model with a Bayes Factor (M1/M0)>100. The logarithmic parameter estimates decreased by 0.70%, given an increase of 1% of the dependent variable.

**Figure 6:**
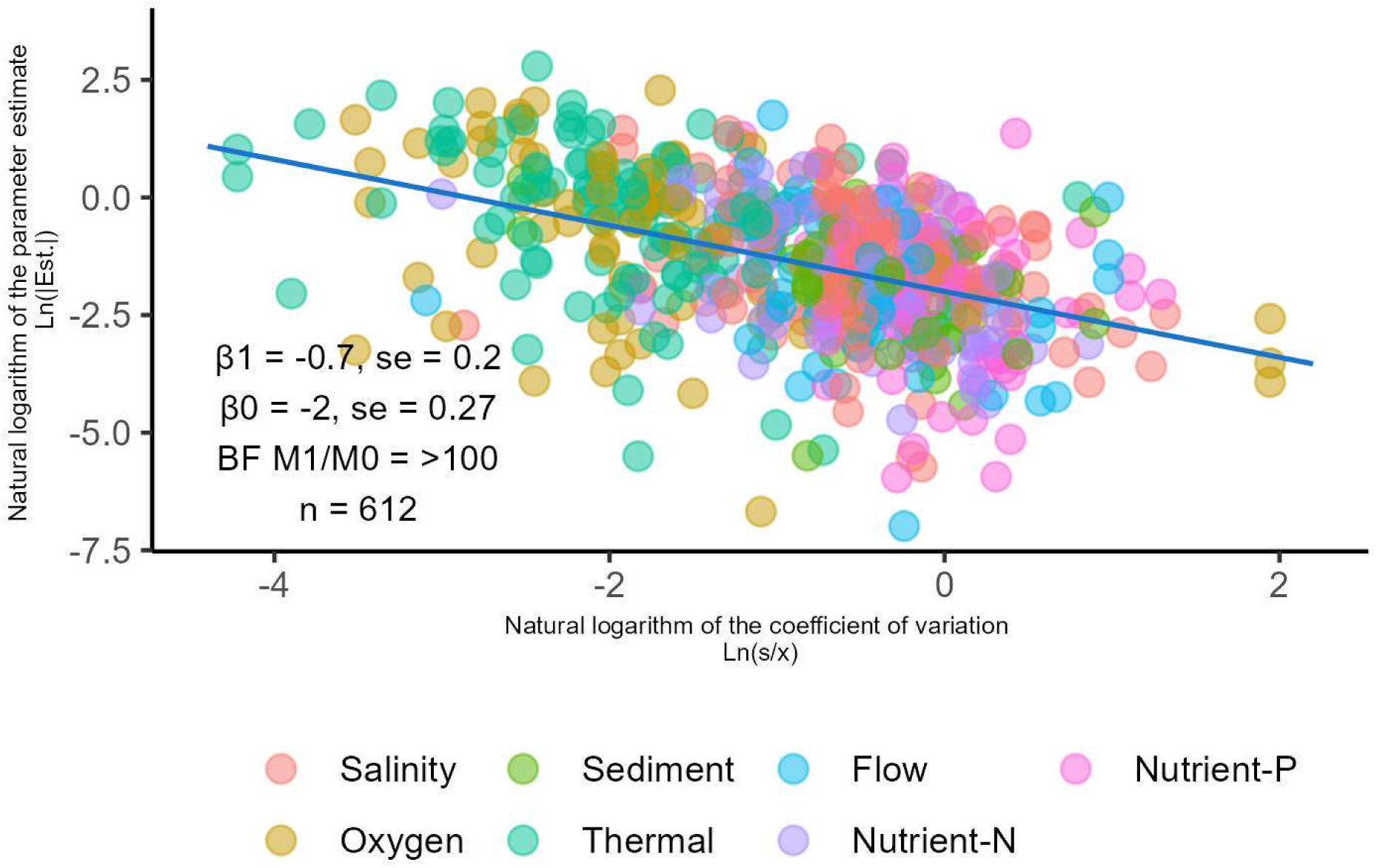
Ln transformed absolute parameter estimates (Ln(|Est.|)) as a function of the coefficient of variation (Ln-transformed).

## 4. Discussion

### 4.1 Stressor response relations across organism groups and stressors

In general, the response variables in the logit-linear model consisted largely of evenness and other proportional metrics, which declined with most stressor gradients. On the other hand, some response variables in log-linear models showed an increase in relation to these stressors (invertebrates and flow). This discrepancy is likely because increasing stressors result in an increase of specific species adapted to such stressor conditions, shifting an evenly distributed community towards a more uneven distribution.

Invertebrates showed the strongest relation to the seven stressors in both logit- and log-models, followed by fish. This is partially due to the larger number of studies available, leading to a better estimation of the stressor-response relation. Related to this, the knowledge of these organism groups is immense, as whole trait databases exist where this group is well represented (Schmidt-Kloiber and Hering, 2015), allowing more targeted investigations that lead to stronger results. This concerns sampling scales, sampling methods for biota and related environmental variables, but also research questions. Finally, invertebrates include many particularly sensitive species responsive to the targeted stressors, while impairments that are rarely addressed might be relevant for microorganisms.

All organism groups are most strongly related to salinity, which is consistent with another study (Kaijser et al., 2024), although bacteria and algae seemed less influenced than macroorganisms. In parts, this might be due to less targeted sampling programs in case of microorganisms, responsive to microscale gradients. Many studies sample from surface water or hard substrates (Kaijser et al., 2024). The taxonomic composition of bacteria or algae may be strongly determined by factors at the microscale, such as the hard substrate minerals, the size of the grains, the concentration of oxygen, bioavailable organic carbon, or electron acceptors in the sediment varying strongly with sediment depth (Smart et al., 2007; Zhang et al., 2020). Moreover, the relationship between microorganisms and salinity is not necessarily indicative of osmotic stress. Salinity is defined by the concentration of all dissolved ions, which causes osmotic stress. However, this does not account for the toxic stresses caused by different ions that are presents in the water. Sea water intrusion can enhance the dominance of sulfate-reducing bacteria because it and increases sulfate concentrations (Kinsman-Costello et al., 2023; Lyman and Fleming, 1940). Yet, this is not the definition of salinity but the direct consequence of the increase in sulfate. For macrophytes, responses are also minor, which could be explained by the small number of studies, considering that macrophytes are strongly determined by salinity under brackish conditions (Kaijser et al., 2019; Remane and Schlieper, 1972).

Lower oxygen concentrations can have severe impacts on invertebrates and fish (Croijmans et al., 2021; Frakes et al., 2021; Jonsson and Jonsson, 2009). For fish activities (e.g., feeding, reproduction, and migration), and egg development is known to be limited by lower oxygen concentrations(Jonsson and Jonsson, 2009; Pauly, 2019; Warren et al., 2015). The dependency of invertebrates on oxygen levels is a textbook example and limits the distribution strongly in case of strong gradients (De Pauw and Vanhooren, 1983; Dodds, 2002); however, nowadays strong oxygen depletion is rarely observed. Photosynthetically active organisms are less sensitive to oxygen depletion, as they generate oxygen.

Fine sediment accumulation was not relatable to algae, bacteria, and archaea. Similar to the explanation above, the diverse biochemical sediment compartment is rarely sampled or included in monitoring programs. In case of invertebrates and fish, fine sediment accumulation might cover relevant proportions of the available habitats (Bartels et al., 2021; Benoy et al., 2012; Louhi et al., 2011). For fish, egg development of characteristic species, such as salmonids, is hampered by sediment deposits (Louhi et al., 2011) and might change food webs (Walker and Walters, 2019).

Higher temperature and lower oxygen concentrations can strongly affect invertebrates and fish (Bonacina et al., 2023; Frakes et al., 2021; Nikinmaa and Rees, 2005). Higher temperature and lower oxygen can also shift the composition of prokaryotic communities towards anaerobic bacteria, resulting in increased production of hydrogen sulfide, which is toxic to most macroorganisms (Kinsman-Costello et al., 2015). This sulfide toxicity interaction has been rarely studied, but may produce additional stress for aquatic organisms (Lamers et al., 2013; Oseid and Smith, 1974; Parveen et al., 2017). Thermal stress had a strong positive relationship with the number of observed fish species, likely due to the higher number of cyprinid species preferring warmer temperature compared to cold-adapted salmonid species (Nikinmaa and Rees, 2005).

Flow alteration and substrate composition are interlinked (Biggs, 1996; Mebane et al., 2014). Flow velocities can have considerable consequences for riverine communities, in particular macroorganisms (Biggs, 1996; Dewson et al., 2007). Macrophyte biomass and community composition are also strongly affected by flow velocity (Chambers et al., 1991; Janauer et al., 2010). There is a scarcity of data on microorganism response to flow. Yet, the heterogeneity of current and flow dynamics within reach-scale also limits interpretation to microscale distributions.

Among the nutrients, algae were strongly related to nitrogen. It is often assumed that the strongest relation would be observed for phosphorus. Macrophytes obtain nutrients primarily from the sediment (Barko et al., 1991; Butcher, 1933; Carignan, 1985) and the relation between surface water nutrients is confounded with dissolved inorganic carbon-related variables (Demars et al., 2012; Kaijser et al., 2021; Maberly et al., 2015). These observations question the suitability of algae and macrophytes as indicators for surface water nutrient stressors in rivers. A direct response of invertebrates and fish to nutrients was not observed. This highlights that more attention is needed to unravel the indirect impacts the total pathways of nutrients in rivers.

Our results underline that microorganisms respond poorly to hydro-morphological stressors (sediment and flow), while these are among the prime stressors for macroorganisms. While salinity is most relevant for all organism groups, oxygen depletion is primarily affecting fish. The response of biota to nutrients is generally weak and almost absent in the case of invertebrates and fish.

### 4.2 Relation of stressor-responses to stressor gradients

We observed a clear relationship between gradient coverage and parameter estimates. This phenomenon has previously been observed (Kaijser et al., 2024; Mack et al., 2022) and can be rated an “statistical artifact” (see Text S2 for a formal explanation). In short, small sample sizes and non-representative samples of only the lower (or higher) part of the environmental gradient result in a low 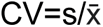 (Fig. 7A). Under these conditions, the model overestimates the model parameter. This is represented in the gray box in Fig. 7C, covering the lower part of the environmental gradient. This gradient section displays a high change per unit increase of the stressor gradient. The higher part of the gradient - beyond the curvature (dashed black line) - has a smaller change per unit increase. Often, samples taken from the entire gradient have a higher variance relative to sample mean, resulting in a higher CV (Fig. 7B). This is not because there is less of an ‘effect’ beyond the curvature due to gradient coverage, but because the samples are better representative of the stressor gradient (Fig. 7C).

**Figure 7:**
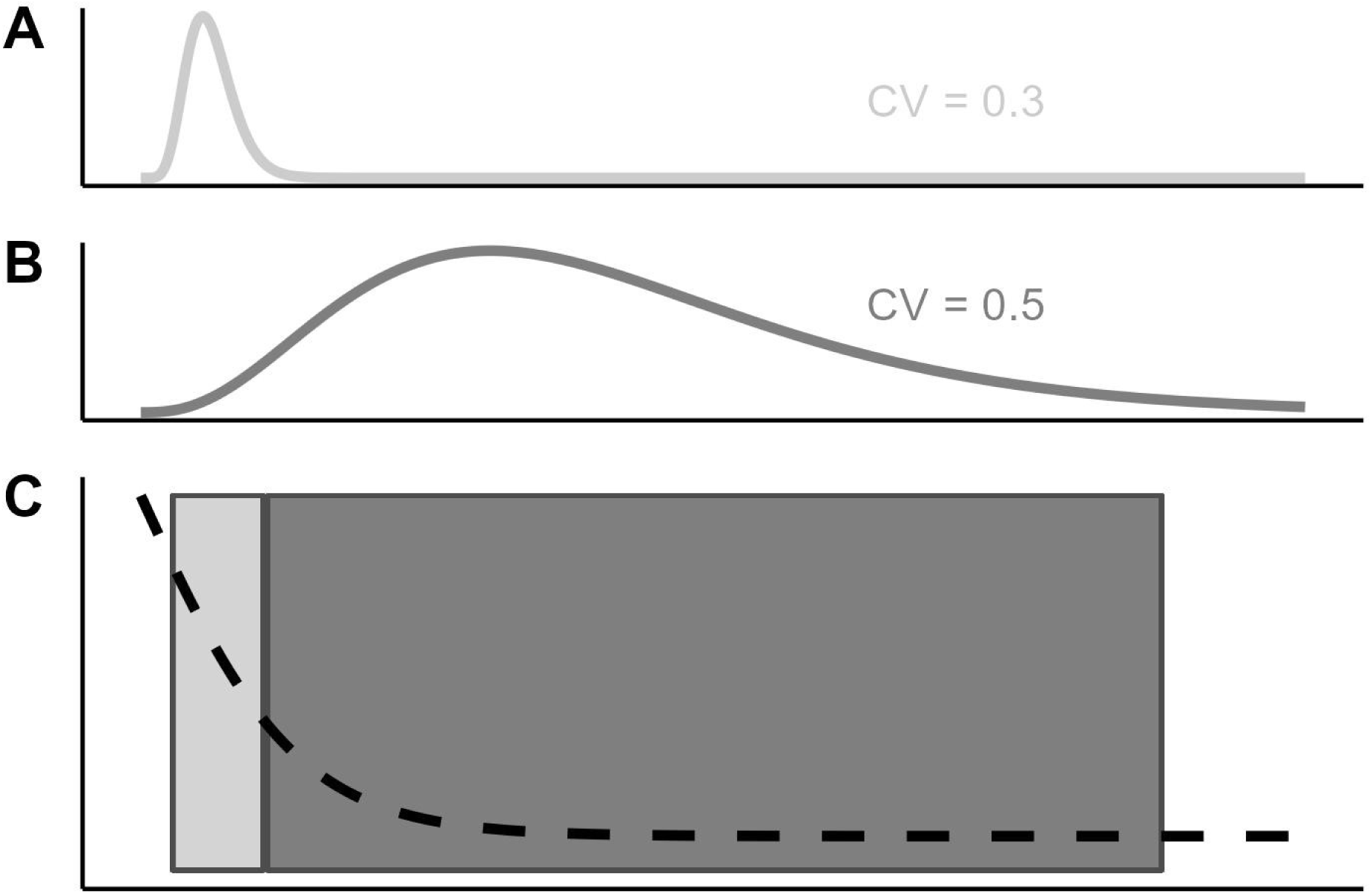
**Visual explanation of the influence of representative sampling of the environmental gradient for a log or logit-linear model.**

The data and results generated can be used as reference material, for re-usage, estimating sample sizes for future studies to prevent overestimation, and prediction, as well as to inform priors for future studies. Furthermore, there exists general consensus among scientists on the directionality (positive or negative) of the given stressor-response relations for different organism groups. However, up to now it has not been attempted to quantitatively summarize the total body of available information on the magnitude of stressor-response relations. Our findings also show that invertebrates were well represented in the data, while other groups were comparatively poorly represented.

## Data availability

Data is available from https://github.com/snwikaij/Data and the functions will be provided under the EcoPostView package on GitHub (after publication).

## Acknowledgements

This paper results from the Collaborative Research Centre 1439 RESIST (Multilevel Response to Stressor Increase and Decrease in Stream Ecosystems; www.sfb-resist.de) funded by the Deutsche Forschungsgemeinschaft (DFG, German Research Foundation; CRC 1439/1, project number: 426547801).

