## Supplementary material for "Global-scale quantification of responses to anthropogenic stressors in six riverine organism groups": Text S1

### Text S1: Methods

#### Data sources and systematic search (Step 1)

A systematic search using various terms in Web-Of-Science (5-3-2024):

phytoplanct* OR macrophyt* OR fish* OR piscine* OR macroinvert* OR benthic Invert* OR invertebrate* OR diatom* OR algae* OR phytobenth* OR bacteria* OR aquatic vegetation* OR aquatic plant* OR submerged vegetation* (Topic) and salini* OR conductiv* OR salt* OR oxygen* OR fine sediment* OR flow* OR velocit* OR discharge* OR temperature* OR thermal* OR nutrient* OR nitrat* OR nitrogen* OR phosph* (Topic) and freshwater OR stream OR river (Topic) not marine* OR lake* OR pond* OR ocean* OR sea* OR terrestrial* OR mammal* OR fungi* OR labor*

This yielded 29,749 articles and after the retrieval phase 22,120 remained (Fig. S1). Each researcher examined 1,000 articles, resulting in a subset from which information was extracted. Before this task an explanation was given in the form of a presentation showing the different biodiversity expression for the different groups (bacteria, algae, macrophytes, invertebrates and fish). These expressions could be number of species, genus, family, order, OTU, community matrices, evenness, shannon-index, or quality metrics (Ecological-Quality-Ratio) or sensitive species (e.g., EPT%). Macrophytes proved to be a challenge and also coverage (%) (macrophytes or bryophytes), was additionally used. These responses were only useful in relation to six stressors: oxygen, salinity, fine sediment, flow, nutrient (nutrient-nitrogen and nutrient-phosphorus) and temperature.

- Oxygen) Oxygen (mg $L^{-1}$),
- Salinity) Conductivity (mS $cm^{-1}$) or CL (mg $L^{-1}$) / 0.6

- Sediment) The fraction of fine sediment or 1-coarse sediments as a proxy for substrate fines.
- Nutrient) Total Nitrogen (mg $L^{-1}$), Nitrate (mg $L^{-1}$), Total Phosphorus (mg $L^{-1}$), phosphate (mg $L^{-1}$), soluble reactive phosphorus (mg $L^{-1}$).
- Flow) Flow velocity (m s) or discharge ($m^3$ $s^1$)
- Thermal) Temperature in degrees Celsius (℃)

A priori, each screener´s performance was assessed on a set of 33 pre-screened articles, some with and some without information that could be extracted from figures, tables or provided datasets. Each participant was informed of their false negatives after which they received their 1,000 articles for screening. The average amount of false negatives was 0.19 (sd=0.15, n=23). False negatives were considered more severe as losing an article is considered worse than obtaining an article that can later be excluded. After this assessment each person screened the 1,000 articles for information that could be extracted from these articles resulting in 3,446 articles remaining.

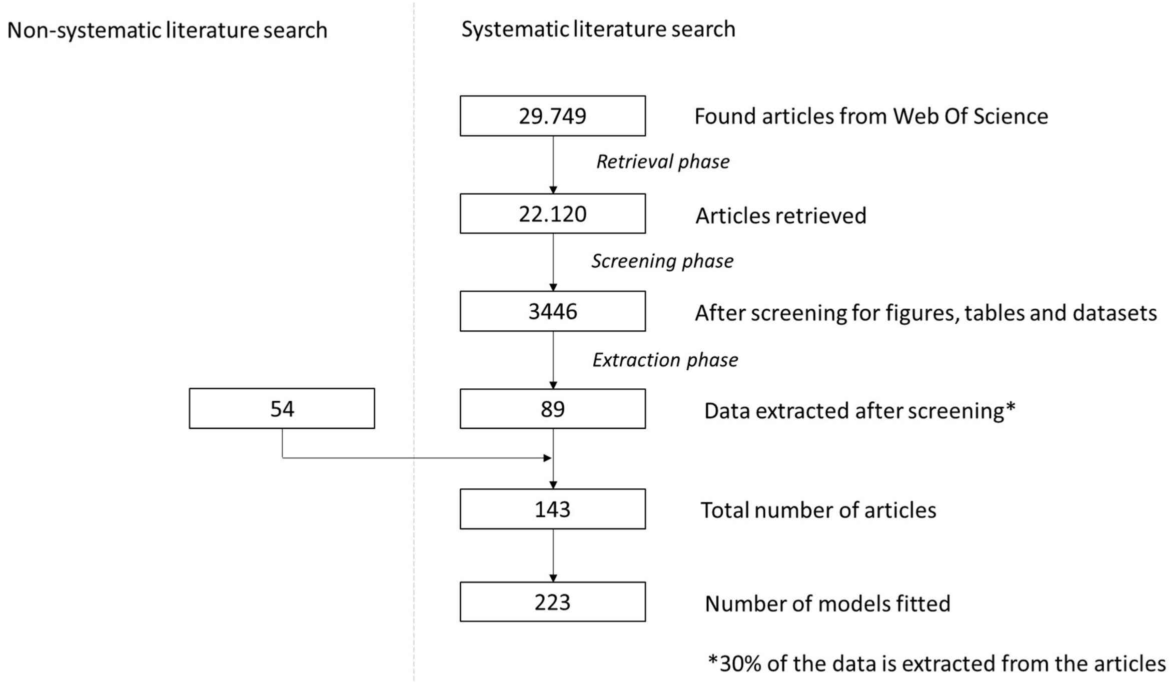

*Figure S1: Representation of a flow chart for the systematic representation of each step selecting the articles.*

During the extraction phase six people extracted the data from the articles. For this we considered only data with an arbitrary amount of six (n>5) or more observations. All appendices that were not directly provided at the end within the paper itself were ignored. From tables and figures presenting the mean and median, the mean was used. If only the median was provided the median was used. If feasible data was extracted from community matrices directly provided in the pdf included tables or appendices. Complex matrices, however, were excluded due to limited resources and time constraints, such as in the directly included appendix of Dolédec et al. (2021). In this example, EPT-taxa% could be extracted. However, the complexity of extracting this from a PDF without mistakes was regarded timewise unfeasible. Evenness was calculated as shannon/Ln(number of taxa) if this was not possible because the matrix, number of species or evenness>1 then evenness=shannon/max(shannon) was used. The full list of all extracted data is provided on GitHub (after publication).

If all data came from figures the type “figure” was appointed to the data, when at least some information originated from a table, it was named “table” and when data was obtained from a dataset, it got the label “dataset”. This was later used in the Egger’s test (step 5). In total 89 articles could be used and were complemented by 54 articles from a non-systematic search.

#### Model fitting (Step 2)

GLM models with log- or logit-link and Negative Binomial or Beta distributed error term were selected (given in the final table). Full models were fitted with up to all seven stressors. The variance of different groups/clustered was separately modeled as random effects if this did not result in convergence issues. If this resulted in a convergence issue a different optimizer was used. If the problem remains, the random effects are dropped.

For each model all independent target variables were transformed using the natural logarithm (Ln) for comparability. Using the two link functions and Ln-transformed gradient the estimated parameter can be easily compared and interpreted as elasticity- or semi-elasticity coefficients (Woolridge, 2001). If the model with log-link is fitted on an Ln-transformed gradient the elasticity coefficient presents the percentage of change in the response relative to one percent change of the stressor. For example, an estimated model parameter of 0.2 would represent a 0.2% increase in the response variable given 1% in the stressor gradient. If a model with logit-link the semi-elasticity coefficient allows for the same interpretation, but precision decreases with distance from 0. This makes it possible to combine different responses i.e., the decline order, family, species or genus relative to different stressor gradients and comparisons between different stressor gradients and organism groups.

For example, if we use a simple linear model where y1={1, 7} and y2={10, 70} are the number of families and species in a sample, respectively, the environmental gradient represents x={25, 350} on a linear scale and the coefficient would be $\beta_1$=(7-1)/(25-350) = -0.185 and $\beta_1$=(70-10)/(25-350) = -0.0185. However, on a Ln-Ln scale the coefficients are exactly the same $\beta_1$=(Ln(7)-Ln(1))/(Ln(25)-Ln(350)) = -0.737 and $\beta_1$=(Ln(70)-Ln(10))/(Ln(25)-Ln(350)) = -0.737. Hence, whether we use family or species level is not directly relevant for the estimation of $\beta_1$, as long as the (levels) families and species approximately follow the same response (e.g., families and species of EPT).

#### Accumulating estimates (Step 3)

Point parameter estimates ($\hat{\beta}$) for intercepts ($\beta_0$) and regression coefficients ($\beta_i$) of the log- and logit-linear models ($g$) with standard errors (se) were extracted from each model: $g\big(E(y|x_n)\big) = \beta_0 + \sum_{i=1}^{n}(\beta_i \cdot Ln(x_i))$. The organism group, stressor type, model type (link function and error distribution), mean and standard deviation of the stressor gradient were all recorded.

#### Prior formulation (Step 4)

The posterior distribution of β can be estimated conditional on prior information using Markov-Chain-Monte-Carlo (MCMC) using the R2JAGS package in R (Su and Yajima, 2021). The MCMC is a simulation method to update our prior information via the likelihood to the posterior probability $\beta_{[stressor\text{-}response,\ link]}$ given the estimations ($\hat{\beta}$). Posterior=Likelihood ∝ Prior:

$$\beta_{[stressor-response,link]} = \{\hat{\beta}_{[stressor-respons\ \ ,link,study]}, \ldots, \hat{\beta}_{[stressor-response,link,study]}\}$$

$$P(\beta_{[stressor-response,link]}|Data, Information)$$
$$= P(Data|\beta_{[stressor-response,link]}) * P(\beta_{[stressor-response,link]}|Information)$$

The prior expresses information into values that are found to be reasonable for our current understanding of the parameter $\beta_1$ (and $\beta_0$). Hence, apriori we can already assign less weight to values that are deemed unreasonable and otherwise assign more weight to values that are more reasonable. We do not have to decide on a single prior, but can include multiple priors and thereby sources of uncertainty. This is called Bayesian Model Averaging (BMA), see Hinne et al. (2020) and for a more formal introduction, see Hoeting et al., (1999). This type of meta-analysis is described as Robust Bayesian Meta Analysis [RoBMA] (Maier et al., 2023). In short, RoBMA uses multiple priors and addresses the relation of the data to these priors. The priors that conform better to the data are more often used in the posterior. By using RoBMA we incorporated eight priors in total. Each stressor response relation got assigned a mix of different priors and weights.

The compression of information through meta-analysis is used to generalize relationships and uncertainties between biotic groups and stressors by aggregating data from diverse sources. The tradition of meta-analysis emphasizes "standardizing effect-sizes". This standardization can be useful under particular context. More importantly is that many sorts of “differences in means“ and “effect-sizes” can be transformed to some “standardized-effect size” and incorporated under the ideal that it is “standardized”.

However, there has been critique on always utilizing and these standardized effect-sizes (Baguley, 2009; Tukey, 1969). It might lose proper error control and ecological interpretation because the units are lost (Baguley, 2009; Tukey, 1969). Hence, the benefits of standardization also have its disadvantages. First, the standardized-effect sizes lose their units and thereby meaning to any ecological interpretation to the field is lost. Cohen’s d is a standardized effect-size and indicates the ratio between the difference in sample means of a treatment and control group divided by the pooled standard deviation. A value of 1 means that the difference in sample means divided by the pooled standard deviation is 1. But what would it mean that the value is 0.5 one can assed the dogmatic “small”, “medium” and “large” terminology, however this was invented by Cohen based on looking at psychological studies and assessing their commonality. Ecology is not psychology and moreover, the interpretations change over time and are relative, rather than universal “truths”. Second, these standardized effect-sizes do not offer an ecologically meaningful interpretation. The standardized effect-size has no unit and is therefore without “sense” and context relative to the field. Assuming that under highly neutral environmental conditions the number of EPT species decreases by 1 compared to a control (MD) and the pooled standard deviation is also one (SD) then Cohen’s d MD/SD=1. Hence, what did we learn about the absolute magnitude an impact of the stressor knowing 1; and can we assume no perturbations of the data were present. Therefore, we focus on the elasticity coefficient as highlighted in step 2.

**Prior formulation**

The relation between chlorophyll-a and total phosphorus is strong and generally believed to represent a causal relation in aquatic ecology. This relation is often regressed with a simple linear model where both dependent and independent variables are log10 or Ln tranformed, i.e., $Ln(y)=\beta_0+\beta_1*Ln(x)$. This is favorable for this analysis as we can interpret and generalize from it considering this analysis. In the case of $Ln(y)=\beta_0+\beta_1*Ln(x)$ (Ln~Ln) for chlorophyll-a/total phosphorus we observe values ~1-1.5 as Ln(chlorophyll-a [ug $L^{-1}$])/Ln(total phosphorus[mg $L^{-1}$]), see for example both Champion and Currie (2000) and Phillips et al. (2008). In this Ln~Ln, values of 1-1.5 represent a 1-1.5% increase in the response with 1% in the stressor gradient unit given all other variables are held constant. For a Logit~Ln expression (semi-elasticity coefficient) is a similar comparison for small values than approximate 5.

Based on this general understanding of these chlorophyll-a and total phosphorus relation we can already exclude values > ~1-1.5 as being unlikely in the prior. Hence, the former relation is a well-known and researched relation and for all stressor-response relations only smaller values are expected. Furthermore, Lorenz et al. (2023) also show values centered around 0. And while not Ln-transformed (but square-root), the magnitude rarely exceeds -1 or 1. Another useful relation is the decline in macrophyte species with an increase in chlorophyll-a (Phillips et al., 1978). It is widely believed to be a causal and relatively strong relationship (no light no macrophyte). Hence, it is generally accepted that high chlorophyll-a prohibits the growth of macrophytes referred to as “regime shifts”. In Kaijser et al. (2024a) only a decline of approximately -0.3 for the log-linear model is observed. This also included multiple other studies that were used to inform the prior. Thus, a stressor-response analyzed in the article falls at least between -1/1.5 and 1/1.5 and has or has a comparable strong relation of approximately -0.3 or 0.3. Comparable, because it seems that experts expect strong stressor-response relation to a specific organism group (e.g., oxygen to invertebrates, but not to algae or macrophytes). However, in Kaijser et al., (2024b) the average of all responses is close to 0, but the estimates for conductivity often center around 0.2. Nonetheless, Kaijser et al., (2024b) did not use an absolute response but a community metric. The advantage is that Kaijser et al., (2024b) also used the Logit~Ln transformation. The idea of a strong or weak relation (-0.3 and 0.3 or -0.2 and 0.2) seems unlikely based on debates around some of the stressor-response relations (e.g., Demars et al., 2012; Demars and Edwards, 2009; Demars and Harper, 1998). If we would pool all estimates from Kaijser et al. (2024a), Lorenz et al. (2023) and Mack et al. (2022) then the average approaches 0 (NULL) association. Therefore, there is a reasonable to incorporate shrinkage centering around 0. Since we applied Bayesian Model Averaging different priors and weights were assigned to different possibilities.

- **Prior (1): Strong alternative**

  N(0.3, 0.1), N(-0.3, 0.1) or N(0, 0.1) depending on the directionality of the assumed association, weighted 12.5%.

  This prior model states the main hypothesis that macro-organism (fish and invertebrates) are often not related to nutrients [N(0, 0.1)]. While macro-organisms such as bacteria and algae are strongly positively related to stressors, such as Nutrient-N (N(0,3, 0.1) and macrophytes are strongly negatively related to Nutrient-N (N(-0.3, 0.1)). The word “strong” is unfortunate as N(0, 0.1) is similar to the null prior (7). However, all priors have been given presente in Tab S1, below and this N(0, 0.1) is later countered by incorporating a prior that would correct if we were wrong about there not being a relation.

- **Prior (2): Alternative weak**

  N(0.2, 0.1), N(-0.2, 0.1) or N(0, 0.1) depending on the directionality of the assumed association, weighted 20%.

This prior indicates the main prior hypothesis, it also receives the heaviest weight (20%). It has the same directionality as the priors formulated above. For example, if the alternative strong prior above is given N(-0.3, 0.1) then this prior is N(-0.2, 0.1). There is an exception, as when no relation is expected, then N(0, 0.1) remains similar.

- **Prior (3): Alternative weak and broad**

N(0.2, 0.2), N(-0.2, 0.2) or N(-0.2, 0.2) depending on the directionality of the assumed association, weighted 12.5%.

This prior has the same average mean as the previous prior. However, the associated relation is assumed to be more scattered, making exactly pinpointing the parameter unclear.

- **Prior (4): Opposing alternative weak and broad**

If we were wrong about the directionality of prior 3 we oppose a “counter” prior, N(0.2, 0.2), N(-0.2, 0.2) or N(-0.2, 0.2) or  depending on the directionality of the assumed association, weighted 5%.

We can of course be wrong about the negative or positive directionality of the estimate as proposed in prior 3. To include the possibility of being wrong an opposing prior is formulated.  This means if prior 3 is N(-0.2, 0.2) this prior proposes the possibility it is actually N(-0.2, 0.2).

- **Prior (5): Opposing strong positive alternative prior for non-relation**

For some estimates no stressor-response relations (Priors 1-3) are expected. However, we might be wrong and therefore only for these relations a positive relation was introduced N(0.3, 0.1) and for the others N(0, 0.1), weighted 12.5%

The hypothesis is that micro-organisms respond not or only weakly to hydro-morphological variables (e.g., *b1_Flow_log_Bacteria*, see Tab S1*).* This means that means of the priors 1-4 are 0. However, also here we might be wrong. This means that for the priors which have a mean in the priors 1-4, we pose a strong positive alternative prior if there is no stressor-response relation expected.

- **Prior (6): Opposing strong negative alternative prior for non-relation**

As to the previous, we might be wrong and therefore only for these relations a negative relation was introduced N(-0.3, 0.1) and for the others N(0, 0.1), weighted 12.5%.

Alternative to prior (5) we might be wrong about priors 1-4 (non stressor-response relation) and wrong about the positive direction of prior 5 and therefore we pose a strong negative alternative prior if there is no stressor-response relation expected.

**Prior (7): Null prior**

Assuming no relation, N(0, 0.1), weighted 12.5%

Clearly it might be possible we can be all wrong and there is simply no stressor-response relation. Therefore, for all stressor-response relations an null prior is introduced.

- **Prior 8) Ignorant prior:**

Assuming no relation, but excluding values <-1.5 and >1.5 as 99% N(0, 0.5), weight 12.5%

This prior excludes extreme values >1.5%. It remains, however, completely “ignorant” about the direction of the estimation.

In the table below the different priors for ($\beta_1$) are given, plus the priors set for each intercept ($\beta_0$) which are later used to create the HOP-lines.

*Table S1: Different priors for each stressor-response relation with their weights.*

| ***Prior*** | ***1*** | | ***2*** | | ***3*** | | ***4*** | | ***5*** | | ***6*** | | ***7*** | | ***8*** | |
|---|---|---|---|---|---|---|---|---|---|---|---|---|---|---|---|---|
| ***Prior weight (prior probability)*** | ***12.5%*** | | ***20%*** | | ***12.5%*** | | ***5%*** | | ***12.5%*** | | ***12.5%*** | | ***12.5%*** | | ***12.5%*** | |
| ***Levels*** | ***mu_1*** | ***se_1*** | ***mu_2*** | ***se_2*** | ***mu_3*** | ***se_3*** | ***mu_4*** | ***se_4*** | ***mu_5*** | ***se_5*** | ***mu_6*** | ***se_6*** | ***mu_7*** | ***se_7*** | ***mu_8*** | ***se_8*** |
| *b0_NA_log_Algae* | *0* | *10* | *0* | *10* | *0* | *10* | *0* | *10* | *0* | *10* | *0* | *10* | *0* | *10* | *0* | *10* |
| *b0_NA_log_Bacteria* | *0* | *10* | *0* | *10* | *0* | *10* | *0* | *10* | *0* | *10* | *0* | *10* | *0* | *10* | *0* | *10* |
| *b0_NA_log_Fish* | *0* | *10* | *0* | *10* | *0* | *10* | *0* | *10* | *0* | *10* | *0* | *10* | *0* | *10* | *0* | *10* |
| *b0_NA_log_Invertebrates* | *0* | *10* | *0* | *10* | *0* | *10* | *0* | *10* | *0* | *10* | *0* | *10* | *0* | *10* | *0* | *10* |
| *b0_NA_log_Macrophytes* | *0* | *10* | *0* | *10* | *0* | *10* | *0* | *10* | *0* | *10* | *0* | *10* | *0* | *10* | *0* | *10* |

| ***Prior*** | ***1*** | | ***2*** | | ***3*** | | ***4*** | | ***5*** | | ***6*** | | ***7*** | | ***8*** | |
|---|---|---|---|---|---|---|---|---|---|---|---|---|---|---|---|---|
| ***Prior weight (prior probability)*** | ***12.5%*** | | ***20%*** | | ***12.5%*** | | ***5%*** | | ***12.5%*** | | ***12.5%*** | | ***12.5%*** | | ***12.5%*** | |
| ***Levels*** | ***mu_1*** | ***se_1*** | ***mu_2*** | ***se_2*** | ***mu_3*** | ***se_3*** | ***mu_4*** | ***se_4*** | ***mu_5*** | ***se_5*** | ***mu_6*** | ***se_6*** | ***mu_7*** | ***se_7*** | ***mu_8*** | ***se_8*** |
| *b0_NA_logit_Algae* | *0* | *10* | *0* | *10* | *0* | *10* | *0* | *10* | *0* | *10* | *0* | *10* | *0* | *10* | *0* | *10* |
| *b0_NA_logit_Bacteria* | *0* | *10* | *0* | *10* | *0* | *10* | *0* | *10* | *0* | *10* | *0* | *10* | *0* | *10* | *0* | *10* |
| *b0_NA_logit_Fish* | *0* | *10* | *0* | *10* | *0* | *10* | *0* | *10* | *0* | *10* | *0* | *10* | *0* | *10* | *0* | *10* |
| *b0_NA_logit_Invertebrates* | *0* | *10* | *0* | *10* | *0* | *10* | *0* | *10* | *0* | *10* | *0* | *10* | *0* | *10* | *0* | *10* |
| *b0_NA_logit_Macrophytes* | *0* | *10* | *0* | *10* | *0* | *10* | *0* | *10* | *0* | *10* | *0* | *10* | *0* | *10* | *0* | *10* |
| *b1_Flow_log_Algae* | *0* | *0.1* | *0* | *0.1* | *0* | *0.2* | *0* | *0.2* | *0* | *0.1* | *0* | *0.1* | *0* | *0.1* | *0* | *0.5* |

| Prior | 1 | | 2 | | 3 | | 4 | | 5 | | 6 | | 7 | | 8 | |
|---|---|---|---|---|---|---|---|---|---|---|---|---|---|---|---|---|
| Prior weight (prior probability) | 12.5% | | 20% | | 12.5% | | 5% | | 12.5% | | 12.5% | | 12.5% | | 12.5% | |
| Levels | mu_1 | se_1 | mu_2 | se_2 | mu_3 | se_3 | mu_4 | se_4 | mu_5 | se_5 | mu_6 | se_6 | mu_7 | se_7 | mu_8 | se_8 |
| b1_Flow_log_Bacteria | 0 | 0.1 | 0 | 0.1 | 0 | 0.2 | 0 | 0.2 | 0 | 0.1 | 0 | 0.1 | 0 | 0.1 | 0 | 0.5 |
| b1_Flow_log_Fish | 0.3 | 0.1 | 0.2 | 0.1 | 0.2 | 0.2 | -0.2 | 0.2 | 0 | 0.1 | 0 | 0.1 | 0 | 0.1 | 0 | 0.5 |
| b1_Flow_log_Invertebrates | 0.3 | 0.1 | 0.2 | 0.1 | 0.2 | 0.2 | -0.2 | 0.2 | 0 | 0.1 | 0 | 0.1 | 0 | 0.1 | 0 | 0.5 |
| b1_Flow_log_Macrophytes | -0.3 | 0.1 | -0.2 | 0.1 | -0.2 | 0.2 | 0.2 | 0.2 | 0 | 0.1 | 0 | 0.1 | 0 | 0.1 | 0 | 0.5 |
| b1_Flow_logit_Fish | -0.3 | 0.1 | -0.2 | 0.1 | -0.2 | 0.2 | 0.2 | 0.2 | 0 | 0.1 | 0 | 0.1 | 0 | 0.1 | 0 | 0.5 |
| b1_Flow_logit_Invertebrates | -0.3 | 0.1 | -0.2 | 0.1 | -0.2 | 0.2 | 0.2 | 0.2 | 0 | 0.1 | 0 | 0.1 | 0 | 0.1 | 0 | 0.5 |

| ***Prior*** | ***1*** | | ***2*** | | ***3*** | | ***4*** | | ***5*** | | ***6*** | | ***7*** | | ***8*** | |
|---|---|---|---|---|---|---|---|---|---|---|---|---|---|---|---|---|
| ***Prior weight (prior probability)*** | ***12.5%*** | | ***20%*** | | ***12.5%*** | | ***5%*** | | ***12.5%*** | | ***12.5%*** | | ***12.5%*** | | ***12.5%*** | |
| ***Levels*** | ***mu_1*** | ***se_1*** | ***mu_2*** | ***se_2*** | ***mu_3*** | ***se_3*** | ***mu_4*** | ***se_4*** | ***mu_5*** | ***se_5*** | ***mu_6*** | ***se_6*** | ***mu_7*** | ***se_7*** | ***mu_8*** | ***se_8*** |
| *b1_Flow_logit_Macrophytes* | *-0.3* | *0.1* | *-0.2* | *0.1* | *-0.2* | *0.2* | *0.2* | *0.2* | *0* | *0.1* | *0* | *0.1* | *0* | *0.1* | *0* | *0.5* |
| *b1_Nutrient-N_log_Algae* | *0.3* | *0.1* | *0.2* | *0.1* | *0.2* | *0.2* | *-0.2* | *0.2* | *0* | *0.1* | *0* | *0.1* | *0* | *0.1* | *0* | *0.5* |
| *b1_Nutrient-N_log_Bacteria* | *0.3* | *0.1* | *0.2* | *0.1* | *0.2* | *0.2* | *-0.2* | *0.2* | *0* | *0.1* | *0* | *0.1* | *0* | *0.1* | *0* | *0.5* |
| *b1_Nutrient-N_log_Fish* | *0* | *0.1* | *0* | *0.1* | *0* | *0.2* | *0* | *0.2* | *0.3* | *0.1* | *-0.3* | *0.1* | *0* | *0.1* | *0* | *0.5* |
| *b1_Nutrient-N_log_Invertebrates* | *0* | *0.1* | *0* | *0.1* | *0* | *0.2* | *0* | *0.2* | *0.3* | *0.1* | *-0.3* | *0.1* | *0* | *0.1* | *0* | *0.5* |

| Prior | 1 | | 2 | | 3 | | 4 | | 5 | | 6 | | 7 | | 8 | |
|---|---|---|---|---|---|---|---|---|---|---|---|---|---|---|---|---|
| Prior weight (prior probability) | 12.5% | | 20% | | 12.5% | | 5% | | 12.5% | | 12.5% | | 12.5% | | 12.5% | |
| Levels | mu_1 | se_1 | mu_2 | se_2 | mu_3 | se_3 | mu_4 | se_4 | mu_5 | se_5 | mu_6 | se_6 | mu_7 | se_7 | mu_8 | se_8 |
| b1_Nutrient-N_log_Macrophytes | -0.3 | 0.1 | -0.2 | 0.1 | -0.2 | 0.2 | 0.2 | 0.2 | 0 | 0.1 | 0 | 0.1 | 0 | 0.1 | 0 | 0.5 |
| b1_Nutrient-N_logit_Algae | 0.3 | 0.1 | 0.2 | 0.1 | 0.2 | 0.2 | -0.2 | 0.2 | 0 | 0.1 | 0 | 0.1 | 0 | 0.1 | 0 | 0.5 |
| b1_Nutrient-N_logit_Bacteria | 0 | 0.1 | 0 | 0.1 | 0 | 0.2 | 0 | 0.2 | 0.3 | 0.1 | -0.3 | 0.1 | 0 | 0.1 | 0 | 0.5 |
| b1_Nutrient-N_logit_Fish | 0 | 0.1 | 0 | 0.1 | 0 | 0.2 | 0 | 0.2 | 0.3 | 0.1 | -0.3 | 0.1 | 0 | 0.1 | 0 | 0.5 |
| b1_Nutrient-N_logit_Invertebrates | 0 | 0.1 | 0 | 0.1 | 0 | 0.2 | 0 | 0.2 | 0.3 | 0.1 | -0.3 | 0.1 | 0 | 0.1 | 0 | 0.5 |

| ***Prior*** | ***1*** | | ***2*** | | ***3*** | | ***4*** | | ***5*** | | ***6*** | | ***7*** | | ***8*** | |
|---|---|---|---|---|---|---|---|---|---|---|---|---|---|---|---|---|
| ***Prior weight (prior probability)*** | ***12.5%*** | | ***20%*** | | ***12.5%*** | | ***5%*** | | ***12.5%*** | | ***12.5%*** | | ***12.5%*** | | ***12.5%*** | |
| ***Levels*** | ***mu_1*** | ***se_1*** | ***mu_2*** | ***se_2*** | ***mu_3*** | ***se_3*** | ***mu_4*** | ***se_4*** | ***mu_5*** | ***se_5*** | ***mu_6*** | ***se_6*** | ***mu_7*** | ***se_7*** | ***mu_8*** | ***se_8*** |
| *b1_Nutrient-N_logit_Macrophytes* | *-0.3* | *0.1* | *-0.2* | *0.1* | *-0.2* | *0.2* | *0.2* | *0.2* | *0* | *0.1* | *0* | *0.1* | *0* | *0.1* | *0* | *0.5* |
| *b1_Nutrient-P_log_Algae* | *0.3* | *0.1* | *0.2* | *0.1* | *0.2* | *0.2* | *-0.2* | *0.2* | *0* | *0.1* | *0* | *0.1* | *0* | *0.1* | *0* | *0.5* |
| *b1_Nutrient-P_log_Bacteria* | *0.3* | *0.1* | *0.2* | *0.1* | *0.2* | *0.2* | *-0.2* | *0.2* | *0* | *0.1* | *0* | *0.1* | *0* | *0.1* | *0* | *0.5* |
| *b1_Nutrient-P_log_Fish* | *0* | *0.1* | *0* | *0.1* | *0* | *0.2* | *0* | *0.2* | *0.3* | *0.1* | *-0.3* | *0.1* | *0* | *0.1* | *0* | *0.5* |
| *b1_Nutrient-P_log_Invertebrates* | *0* | *0.1* | *0* | *0.1* | *0* | *0.2* | *0* | *0.2* | *0.3* | *0.1* | *-0.3* | *0.1* | *0* | *0.1* | *0* | *0.5* |

| **Prior** | **1** | | **2** | | **3** | | **4** | | **5** | | **6** | | **7** | | **8** | |
|---|---|---|---|---|---|---|---|---|---|---|---|---|---|---|---|---|
| **Prior weight (prior probability)** | **12.5%** | | **20%** | | **12.5%** | | **5%** | | **12.5%** | | **12.5%** | | **12.5%** | | **12.5%** | |
| **Levels** | **mu_1** | **se_1** | **mu_2** | **se_2** | **mu_3** | **se_3** | **mu_4** | **se_4** | **mu_5** | **se_5** | **mu_6** | **se_6** | **mu_7** | **se_7** | **mu_8** | **se_8** |
| b1_Nutrient-P_log_Macrophytes | -0.3 | 0.1 | -0.2 | 0.1 | -0.2 | 0.2 | 0.2 | 0.2 | 0 | 0.1 | 0 | 0.1 | 0 | 0.1 | 0 | 0.5 |
| b1_Nutrient-P_logit_Algae | 0.3 | 0.1 | 0.2 | 0.1 | 0.2 | 0.2 | -0.2 | 0.2 | 0 | 0.1 | 0 | 0.1 | 0 | 0.1 | 0 | 0.5 |
| b1_Nutrient-P_logit_Bacteria | 0 | 0.1 | 0 | 0.1 | 0 | 0.2 | 0 | 0.2 | 0.3 | 0.1 | -0.3 | 0.1 | 0 | 0.1 | 0 | 0.5 |
| b1_Nutrient-P_logit_Fish | 0 | 0.1 | 0 | 0.1 | 0 | 0.2 | 0 | 0.2 | 0.3 | 0.1 | -0.3 | 0.1 | 0 | 0.1 | 0 | 0.5 |
| b1_Nutrient-P_logit_Invertebrates | 0 | 0.1 | 0 | 0.1 | 0 | 0.2 | 0 | 0.2 | 0.3 | 0.1 | -0.3 | 0.1 | 0 | 0.1 | 0 | 0.5 |

| ***Prior*** | ***1*** | | ***2*** | | ***3*** | | ***4*** | | ***5*** | | ***6*** | | ***7*** | | ***8*** | |
|---|---|---|---|---|---|---|---|---|---|---|---|---|---|---|---|---|
| ***Prior weight (prior probability)*** | ***12.5%*** | | ***20%*** | | ***12.5%*** | | ***5%*** | | ***12.5%*** | | ***12.5%*** | | ***12.5%*** | | ***12.5%*** | |
| ***Levels*** | ***mu_1*** | ***se_1*** | ***mu_2*** | ***se_2*** | ***mu_3*** | ***se_3*** | ***mu_4*** | ***se_4*** | ***mu_5*** | ***se_5*** | ***mu_6*** | ***se_6*** | ***mu_7*** | ***se_7*** | ***mu_8*** | ***se_8*** |
| *b1_Nutrient-P_logit_Macrophytes* | *-0.3* | *0.1* | *-0.2* | *0.1* | *-0.2* | *0.2* | *0.2* | *0.2* | *0* | *0.1* | *0* | *0.1* | *0* | *0.1* | *0* | *0.5* |
| *b1_Oxygen_log_Algae* | *0* | *0.1* | *0* | *0.1* | *0* | *0.2* | *0* | *0.2* | *0.3* | *0.1* | *-0.3* | *0.1* | *0* | *0.1* | *0* | *0.5* |
| *b1_Oxygen_log_Bacteria* | *0.3* | *0.1* | *0.2* | *0.1* | *0.2* | *0.2* | *-0.2* | *0.2* | *0* | *0.1* | *0* | *0.1* | *0* | *0.1* | *0* | *0.5* |
| *b1_Oxygen_log_Fish* | *0.3* | *0.1* | *0.2* | *0.1* | *0.2* | *0.2* | *-0.2* | *0.2* | *0* | *0.1* | *0* | *0.1* | *0* | *0.1* | *0* | *0.5* |
| *b1_Oxygen_log_Invertebrates* | *0.3* | *0.1* | *0.2* | *0.1* | *0.2* | *0.2* | *-0.2* | *0.2* | *0* | *0.1* | *0* | *0.1* | *0* | *0.1* | *0* | *0.5* |

| **Prior** | **1** | | **2** | | **3** | | **4** | | **5** | | **6** | | **7** | | **8** | |
|---|---|---|---|---|---|---|---|---|---|---|---|---|---|---|---|---|
| **Prior weight (prior probability)** | **12.5%** | | **20%** | | **12.5%** | | **5%** | | **12.5%** | | **12.5%** | | **12.5%** | | **12.5%** | |
| **Levels** | **mu_1** | **se_1** | **mu_2** | **se_2** | **mu_3** | **se_3** | **mu_4** | **se_4** | **mu_5** | **se_5** | **mu_6** | **se_6** | **mu_7** | **se_7** | **mu_8** | **se_8** |
| b1_Oxygen_log_Macrophytes | 0 | 0.1 | 0 | 0.1 | 0 | 0.2 | 0 | 0.2 | 0.3 | 0.1 | -0.3 | 0.1 | 0 | 0.1 | 0 | 0.5 |
| b1_Oxygen_logit_Algae | 0 | 0.1 | 0 | 0.1 | 0 | 0.2 | 0 | 0.2 | 0.3 | 0.1 | -0.3 | 0.1 | 0 | 0.1 | 0 | 0.5 |
| b1_Oxygen_logit_Bacteria | 0 | 0.1 | 0 | 0.1 | 0 | 0.2 | 0 | 0.2 | 0.3 | 0.1 | -0.3 | 0.1 | 0 | 0.1 | 0 | 0.5 |
| b1_Oxygen_logit_Fish | 0.3 | 0.1 | 0.2 | 0.1 | 0.2 | 0.2 | -0.2 | 0.2 | 0 | 0.1 | 0 | 0.1 | 0 | 0.1 | 0 | 0.5 |
| b1_Oxygen_logit_Invertebrates | 0.3 | 0.1 | 0.2 | 0.1 | 0.2 | 0.2 | -0.2 | 0.2 | 0 | 0.1 | 0 | 0.1 | 0 | 0.1 | 0 | 0.5 |
| b1_Oxygen_logit_Macrophytes | 0 | 0.1 | 0 | 0.1 | 0 | 0.2 | 0 | 0.2 | 0.3 | 0.1 | -0.3 | 0.1 | 0 | 0.1 | 0 | 0.5 |

| **Prior** | **1** | | **2** | | **3** | | **4** | | **5** | | **6** | | **7** | | **8** | |
|---|---|---|---|---|---|---|---|---|---|---|---|---|---|---|---|---|
| **Prior weight (prior probability)** | **12.5%** | | **20%** | | **12.5%** | | **5%** | | **12.5%** | | **12.5%** | | **12.5%** | | **12.5%** | |
| **Levels** | **mu_1** | **se_1** | **mu_2** | **se_2** | **mu_3** | **se_3** | **mu_4** | **se_4** | **mu_5** | **se_5** | **mu_6** | **se_6** | **mu_7** | **se_7** | **mu_8** | **se_8** |
| *b1_Salinity_log_Algae* | *-0.3* | *0.1* | *-0.2* | *0.1* | *-0.2* | *0.2* | *0.2* | *0.2* | *0* | *0.1* | *0* | *0.1* | *0* | *0.1* | *0* | *0.5* |
| *b1_Salinity_log_Bacteria* | *-0.3* | *0.1* | *-0.2* | *0.1* | *-0.2* | *0.2* | *0.2* | *0.2* | *0* | *0.1* | *0* | *0.1* | *0* | *0.1* | *0* | *0.5* |
| *b1_Salinity_log_Fish* | *-0.3* | *0.1* | *-0.2* | *0.1* | *-0.2* | *0.2* | *0.2* | *0.2* | *0* | *0.1* | *0* | *0.1* | *0* | *0.1* | *0* | *0.5* |
| *b1_Salinity_log_Invertebrates* | *-0.3* | *0.1* | *-0.2* | *0.1* | *-0.2* | *0.2* | *0.2* | *0.2* | *0* | *0.1* | *0* | *0.1* | *0* | *0.1* | *0* | *0.5* |
| *b1_Salinity_log_Macrophytes* | *-0.3* | *0.1* | *-0.2* | *0.1* | *-0.2* | *0.2* | *0.2* | *0.2* | *0* | *0.1* | *0* | *0.1* | *0* | *0.1* | *0* | *0.5* |
| *b1_Salinity_logit_Algae* | *-0.3* | *0.1* | *-0.2* | *0.1* | *-0.2* | *0.2* | *0.2* | *0.2* | *0* | *0.1* | *0* | *0.1* | *0* | *0.1* | *0* | *0.5* |

| ***Prior*** | ***1*** | | ***2*** | | ***3*** | | ***4*** | | ***5*** | | ***6*** | | ***7*** | | ***8*** | |
|---|---|---|---|---|---|---|---|---|---|---|---|---|---|---|---|---|
| ***Prior weight (prior probability)*** | ***12.5%*** | | ***20%*** | | ***12.5%*** | | ***5%*** | | ***12.5%*** | | ***12.5%*** | | ***12.5%*** | | ***12.5%*** | |
| ***Levels*** | ***mu_1*** | ***se_1*** | ***mu_2*** | ***se_2*** | ***mu_3*** | ***se_3*** | ***mu_4*** | ***se_4*** | ***mu_5*** | ***se_5*** | ***mu_6*** | ***se_6*** | ***mu_7*** | ***se_7*** | ***mu_8*** | ***se_8*** |
| *b1_Salinity_logit_Bacteria* | *-0.3* | *0.1* | *-0.2* | *0.1* | *-0.2* | *0.2* | *0.2* | *0.2* | *0* | *0.1* | *0* | *0.1* | *0* | *0.1* | *0* | *0.5* |
| *b1_Salinity_logit_Fish* | *-0.3* | *0.1* | *-0.2* | *0.1* | *-0.2* | *0.2* | *0.2* | *0.2* | *0* | *0.1* | *0* | *0.1* | *0* | *0.1* | *0* | *0.5* |
| *b1_Salinity_logit_Invertebrates* | *-0.3* | *0.1* | *-0.2* | *0.1* | *-0.2* | *0.2* | *0.2* | *0.2* | *0* | *0.1* | *0* | *0.1* | *0* | *0.1* | *0* | *0.5* |
| *b1_Salinity_logit_Macrophytes* | *-0.3* | *0.1* | *-0.2* | *0.1* | *-0.2* | *0.2* | *0.2* | *0.2* | *0* | *0.1* | *0* | *0.1* | *0* | *0.1* | *0* | *0.5* |
| *b1_Sediment_log_Algae* | *0* | *0.1* | *0* | *0.1* | *0* | *0.2* | *0* | *0.2* | *0.3* | *0.1* | *-0.3* | *0.1* | *0* | *0.1* | *0* | *0.5* |
| *b1_Sediment_log_Bacteria* | *0* | *0.1* | *0* | *0.1* | *0* | *0.2* | *0* | *0.2* | *0.3* | *0.1* | *-0.3* | *0.1* | *0* | *0.1* | *0* | *0.5* |

| ***Prior*** | ***1*** | | ***2*** | | ***3*** | | ***4*** | | ***5*** | | ***6*** | | ***7*** | | ***8*** | |
|---|---|---|---|---|---|---|---|---|---|---|---|---|---|---|---|---|
| ***Prior weight (prior probability)*** | ***12.5%*** | | ***20%*** | | ***12.5%*** | | ***5%*** | | ***12.5%*** | | ***12.5%*** | | ***12.5%*** | | ***12.5%*** | |
| ***Levels*** | ***mu_1*** | ***se_1*** | ***mu_2*** | ***se_2*** | ***mu_3*** | ***se_3*** | ***mu_4*** | ***se_4*** | ***mu_5*** | ***se_5*** | ***mu_6*** | ***se_6*** | ***mu_7*** | ***se_7*** | ***mu_8*** | ***se_8*** |
| *b1_Sediment_log_Fish* | *-0.3* | *0.1* | *-0.2* | *0.1* | *-0.2* | *0.2* | *0.2* | *0.2* | *0* | *0.1* | *0* | *0.1* | *0* | *0.1* | *0* | *0.5* |
| *b1_Sediment_log_Invertebrates* | *-0.3* | *0.1* | *-0.2* | *0.1* | *-0.2* | *0.2* | *0.2* | *0.2* | *0* | *0.1* | *0* | *0.1* | *0* | *0.1* | *0* | *0.5* |
| *b1_Sediment_log_Macrophytes* | *0* | *0.1* | *0* | *0.1* | *0* | *0.2* | *0* | *0.2* | *0.3* | *0.1* | *-0.3* | *0.1* | *0* | *0.1* | *0* | *0.5* |
| *b1_Sediment_logit_Algae* | *0* | *0.1* | *0* | *0.1* | *0* | *0.2* | *0* | *0.2* | *0.3* | *0.1* | *-0.3* | *0.1* | *0* | *0.1* | *0* | *0.5* |
| *b1_Sediment_logit_Bacteria* | *0* | *0.1* | *0* | *0.1* | *0* | *0.2* | *0* | *0.2* | *0.3* | *0.1* | *-0.3* | *0.1* | *0* | *0.1* | *0* | *0.5* |
| *b1_Sediment_logit_Fish* | *0* | *0.1* | *0* | *0.1* | *0* | *0.2* | *0* | *0.2* | *0.3* | *0.1* | *-0.3* | *0.1* | *0* | *0.1* | *0* | *0.5* |

| **Prior** | **1** | | **2** | | **3** | | **4** | | **5** | | **6** | | **7** | | **8** | |
|---|---|---|---|---|---|---|---|---|---|---|---|---|---|---|---|---|
| **Prior weight (prior probability)** | **12.5%** | | **20%** | | **12.5%** | | **5%** | | **12.5%** | | **12.5%** | | **12.5%** | | **12.5%** | |
| **Levels** | **mu_1** | **se_1** | **mu_2** | **se_2** | **mu_3** | **se_3** | **mu_4** | **se_4** | **mu_5** | **se_5** | **mu_6** | **se_6** | **mu_7** | **se_7** | **mu_8** | **se_8** |
| *b1_Sediment_logit_Invertebrates* | *-0.3* | *0.1* | *-0.2* | *0.1* | *-0.2* | *0.2* | *0.2* | *0.2* | *0* | *0.1* | *0* | *0.1* | *0* | *0.1* | *0* | *0.5* |
| *b1_Sediment_logit_Macrophytes* | *0* | *0.1* | *0* | *0.1* | *0* | *0.2* | *0* | *0.2* | *0.3* | *0.1* | *-0.3* | *0.1* | *0* | *0.1* | *0* | *0.5* |
| *b1_Thermal_log_Algae* | *0.3* | *0.1* | *0.2* | *0.1* | *0.2* | *0.2* | *-0.2* | *0.2* | *0* | *0.1* | *0* | *0.1* | *0* | *0.1* | *0* | *0.5* |
| *b1_Thermal_log_Bacteria* | *0.3* | *0.1* | *0.2* | *0.1* | *0.2* | *0.2* | *-0.2* | *0.2* | *0* | *0.1* | *0* | *0.1* | *0* | *0.1* | *0* | *0.5* |
| *b1_Thermal_log_Fish* | *0.3* | *0.1* | *0.2* | *0.1* | *0.2* | *0.2* | *-0.2* | *0.2* | *0* | *0.1* | *0* | *0.1* | *0* | *0.1* | *0* | *0.5* |
| *b1_Thermal_log_Invertebrates* | *-0.3* | *0.1* | *-0.2* | *0.1* | *-0.2* | *0.2* | *0.2* | *0.2* | *0* | *0.1* | *0* | *0.1* | *0* | *0.1* | *0* | *0.5* |

| *Prior* | *1* | | *2* | | *3* | | *4* | | *5* | | *6* | | *7* | | *8* | |
|---|---|---|---|---|---|---|---|---|---|---|---|---|---|---|---|---|
| ***Prior weight (prior probability)*** | ***12.5%*** | | ***20%*** | | ***12.5%*** | | ***5%*** | | ***12.5%*** | | ***12.5%*** | | ***12.5%*** | | ***12.5%*** | |
| ***Levels*** | ***mu_1*** | ***se_1*** | ***mu_2*** | ***se_2*** | ***mu_3*** | ***se_3*** | ***mu_4*** | ***se_4*** | ***mu_5*** | ***se_5*** | ***mu_6*** | ***se_6*** | ***mu_7*** | ***se_7*** | ***mu_8*** | ***se_8*** |
| *b1_Thermal_log_Macrophytes* | *0.3* | *0.1* | *0.2* | *0.1* | *0.2* | *0.2* | *-0.2* | *0.2* | *0* | *0.1* | *0* | *0.1* | *0* | *0.1* | *0* | *0.5* |
| *b1_Thermal_logit_Algae* | *0.3* | *0.1* | *0.2* | *0.1* | *0.2* | *0.2* | *-0.2* | *0.2* | *0* | *0.1* | *0* | *0.1* | *0* | *0.1* | *0* | *0.5* |
| *b1_Thermal_logit_Bacteria* | *0.3* | *0.1* | *0.2* | *0.1* | *0.2* | *0.2* | *-0.2* | *0.2* | *0* | *0.1* | *0* | *0.1* | *0* | *0.1* | *0* | *0.5* |
| *b1_Thermal_logit_Fish* | *0.3* | *0.1* | *0.2* | *0.1* | *0.2* | *0.2* | *-0.2* | *0.2* | *0* | *0.1* | *0* | *0.1* | *0* | *0.1* | *0* | *0.5* |
| *b1_Thermal_logit_Invertebrates* | *-0.3* | *0.1* | *-0.2* | *0.1* | *-0.2* | *0.2* | *0.2* | *0.2* | *0* | *0.1* | *0* | *0.1* | *0* | *0.1* | *0* | *0.5* |
| *b1_Thermal_logit_Macrophytes* | *0.3* | *0.1* | *0.2* | *0.1* | *0.2* | *0.2* | *-0.2* | *0.2* | *0* | *0.1* | *0* | *0.1* | *0* | *0.1* | *0* | *0.5* |

Using RoBMA, we can express the amount of “learning” over all different priors by dividing the weight the prior occurred in the posterior (posterior odds) by its initial prior weight (prior odds) called the posterior/prior-odds-ratio. For example, 0.2/0.2=1, then this can be expressed as a fraction (0.2/0.2)/(1+(0.2/0.2))=0.5 which expresses no information was gained. Otherwise >0.5 indicates information gain in favor of the prior, and <0.5 indicates information gain against a prior. Subtracting 0.5, taking the absolute of this subtracting and multiplying by 2, generates a value between 0 and 1. Here, 0 indicates no information gained by introducing data and no shift from prior to posterior odds occurred. This means introducing data did not change anything about our prior statements. A value closer to 1 means that the posterior favors specific prior model(s) more. It would be rather tedious to look at each of the eight prior models for each biotic group and stressor thus we can express the change in information as the “multi prior posterior information ratio” (mmpir).

$$mppir = \frac{1}{n}\sum_{k=1}^{n} 2 \cdot \left| \frac{\frac{P(\beta_k|Data,Model_k)}{P(Model_k,\beta_k)}}{\left(1+\frac{P(\beta_k|Data,Model_k)}{P(Model_k,\beta_k)}\right)} - 0.5 \right| \quad (Eq.\ S1)$$

This is a relative expression of information and completely dependent on the priors used. It is only useful to compare it for a single stressor-response relation and not between stressor-response relations. Moreover, this has to be interpreted with caution, completely information deprived priors will hardly be selected in the posterior and thus contributed much more to the mppir.

#### Data assessment (Step 5)

We investigated potential biases as highlighted by a relation between the precision (1/standard error) and parameter estimates divided by the standard error (se) (Parameter estimate/se). Regressing Parameter estimate/se onto 1/se provides an indication about potential biases. A strong shift in the intercept from a null model would be indicative for publication bias publishing higher Parameter estimate/se with higher se. This relation was investigated using a Linear Mixed

Model more classical known as the Egger's test. In this model the source (Figure, Table or Dataset) was modeled as a random intercept and nested under this each study. Additionally, the regression coefficient for each source was modeled separately. Two priors (data generating models) were used. The alternative model (M1) of the intercept $\beta_0$ models a potential shift. It is a mixed distribution of two normal distributions (denoted as N(mean, standard deviation) with mean of -0.1 or 0.1 with a standard deviation of 0.1 occurring balanced (50% each) in the prior. The null model (M0) is a single model that models the absence of a shift N(0, 0.1). The prior for the slope $\beta_1$ was left intentionally vague N(0, 1). The standard error of the model was set as uniform ranging between 0 and 10 U(0, 10). The prior odds ratio was set at 1 meaning each model is assumed equally likely. The Bayes Factor (BF) then represents the marginal likelihood ratio of how much the data favors M1 over M0 (M1/M0).

#### Relation between parameter and gradient (Step 6)

We expected the relation between the shift in the scale (variance) and location (mean) parameter to be negatively related to the estimate of the regression models. Hence, at the lower part of the gradient higher estimates are expected because often more sensitive species are present. For this, the Ln(|Est.|) was regressed on the Ln of the sample standard deviation (s) divided by the sample mean of the stressor gradient ($\bar{x}$). This represents the natural logarithm of the coefficient of variation (Ln(s/$\bar{x}$)). The expectation was that this relation would be moderate around 0.25 (or 1-0.75) according to the Common Language Effect Size (CLES). CLES indicates how more often x ranks higher than y where 0.5 indicates 50% (50/50) 1=100% and 0=0%. According to the inverse of Dunlap (1994) a value of 0.25 would be would be around sin($\boldsymbol{\pi}$*(0.25-0.5))=-0.71 on increase of Ln(|Est.|) increase with one unit in Ln(s/$\bar{x}$) on a standardized scale. We fitted an LMM modeling both the slope and intercept for each stressor type as random. The prior was formulated based on and the underlying assumption that the response and stressor follow two standard normals then two times is 0.355 to include 0 at the 97.5% lower edge. The prior for M1 was selected as N(-0.71, 0.355) and for M0 N(0, 0.1). With a prior odds ratio of 2 meaning M1 was weighted by 66% and M0 by 33%.

#### Meta-analysis (Step 7)

Each parameter estimate of stressor-response relation per organism group and link-function were modeled separately (level) and each parameter estimate was modeled separately. Also, the variance (se) was modeled separately for each of these. The parameter estimates were found dependent on an expression of gradient length (step 6.) Therefore, this dependency was included in the model as a fixed effect modeling the pooled estimate as a function of the hierarchy and Ln(s/x̄) additionally adjusting for this dependency. Only a part of the likelihood is given in this pre-print:

For each estimate, i={1,… ,n}:

For each level, level={1,… ,n}:

- Individual estimates:

$$est_i \sim N(\mu 2_i, \tau 2_i)$$

$$\tau 2_i \sim \frac{1}{{se_i}^2}$$

- Mean of each level:

$$\mu 2_i \sim N(\mu 1_i, \tau 1_{level,i})$$

- Mean based on level and gradient:

$$\mu 1_i = \mu 1_{level,i} \quad + Ln(CV)_i * adjust_{level,i}$$

The MCMC-iterations used 25 chains with 50000 iterations thinned by 50. The mixing of the chains was only acceptable when Rhat < 1.01 and effective sample sizes >3000.

#### Visualizing the results (Step 8)

For each pooled parameter estimate $\beta_0$ and $\beta_1$ from the meta-analysis presented above the posterior is obtained via the MCMC iterations. The results of these iterations are displayed as posterior probabilities density of $\beta_1$ and provide the full results in the appendix (Tab S2). Furthermore, we can display the model (regression line) for a few of the strongest relations as observed in the posterior estimates $\beta_1$. Hence, for each stressor gradient a set value can be generated $x=\{x_i, \ldots, x_n\}$ and a random value can be drawn from the posterior distribution of $\beta_0$ and $\beta_1$. Then the Ln(x) can be multiplied with the posterior estimate and the inverse of the link function can be applied to plot E(y|x). Then 1,500 random lines generated from the posterior named Hypothetical-Outcome-Plots or in short HOPs (Kale et al., 2019). Each line displays the expected change along the stressor gradient.

A minor addition to the results left out of the main text is that each prior is a quantitative expression of a specific hypothesis. We assessed the overall performance of all stressor-response relations of these priors by calculating how much the data shifted in favor of these prior hypotheses as as the “posterior/prior odds ratio”. Values >1 mean a favor against the prior model and <1 against. The data favored three priors and associated hypotheses most: The weak alternative prior (3), the null prior (7) and the opposing strong negative alternative prior (6). Where most favored by the data. It thus highlights that most of our stressor-response relations are compatible with the weak alternative or null model.

*Table S2: Posterior/prior-odds-ratios for the total analysis representing information gain over the total study.*

| Prior | 1 | 2 | 3 | 4 | 5 | 6 | 7 | 8 |
|---|---|---|---|---|---|---|---|---|
| Name | Strong alternative | Weak alternative | Weak and broad alternative | Opposing alternative weak and broad prior (to account for being wrong about 1-3) | Opposing strong positive alternative prior (when non-relation was expected at 1-4) | Opposing strong negative alternative prior (when non-relation was expected at 1-4) | Null prior | Ignorant prior |
| Prior probability | 12.5% | 20% | 12.5% | 5% | 12.5% | 12.5% | 12.5% | 12.5% |
| Posterior/prior odds ratio = (Posterior probability/prior probability) | 1.03 | 1.07 | 1.01 | 0.91 | 0.92 | 1.04 | 1.05 | 0.86 |
