## Supplementary material for "Global-scale quantification of responses to anthropogenic stressors in six riverine organism groups": Fig. S2

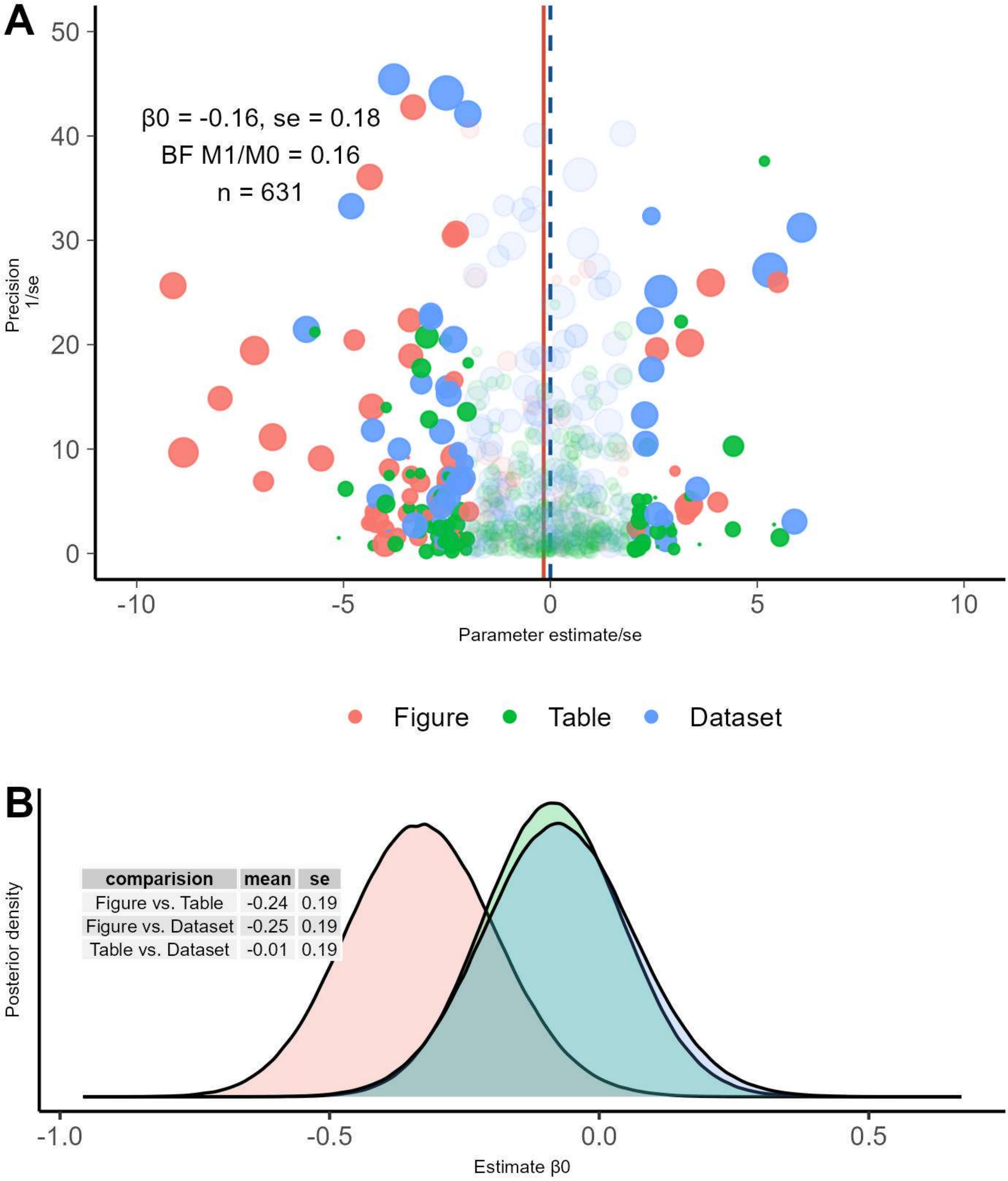


*Figure S2: A) Funnel plot of 1 divided by the standard error (1/se) on the y-axis and parameter divided by the standard error (Parameter estimate/se) on the x-axis. B) Estimates of the difference in means of the intercepts of panel A for each data source in the linear mixed model. The solid red line represents the regression line of the expected value and the dashed blue line the intercept only. Under theoretical ideal conditions both lines should match. The size of the points represents sample size, the color the type of data source (figure, table and dataset) and the transparency the z-score (transparent |Z|<1.96 or p>.05). β0 = intercept, β1 = regression coefficient, BF = Bayes Factor, n = sample size and se = standard error of β1.*
