## Supplementary material for "Global-scale quantification of responses to anthropogenic stressors in six riverine organism groups": Text S2

### Text S2: Formal explanation

Assuming that the underlying stressor gradient has a data generating process following a lower truncated Normal distribution on a log scale (or log normal). The value of the truncation originates from a Gamma distribution and is “stochastic”, the mean is a fixed sequence, the variance is correlated, the sample size taken from stressor gradient is distributed according to a negative binomial distribution with size (r) of and mean (m) and the model fitted to the data is a log-linear model with Poisson error term:

$$\mu = \{i, \ldots, n\} \; and \; \sigma = \mu; \; r = 1, m = 40$$

$$\gamma \sim Gamma(\alpha, \beta)$$

$$n \sim NB(r, m)$$

$$x \sim N(n, Ln(\frac{\mu}{\sqrt{\mu^2/\sigma^2 + 1}}), Ln(\mu^2/\sigma^2 + 1), \gamma, \infty)$$

$$y \sim Poisson(e^{(\beta * Ln(x))})$$

Then due to the noise introduced by the lower truncation of the random variable and smaller sample size (n) the estimate of β is negatively correlated with $Ln(\sqrt{var(x)}/\bar{x}(x))$

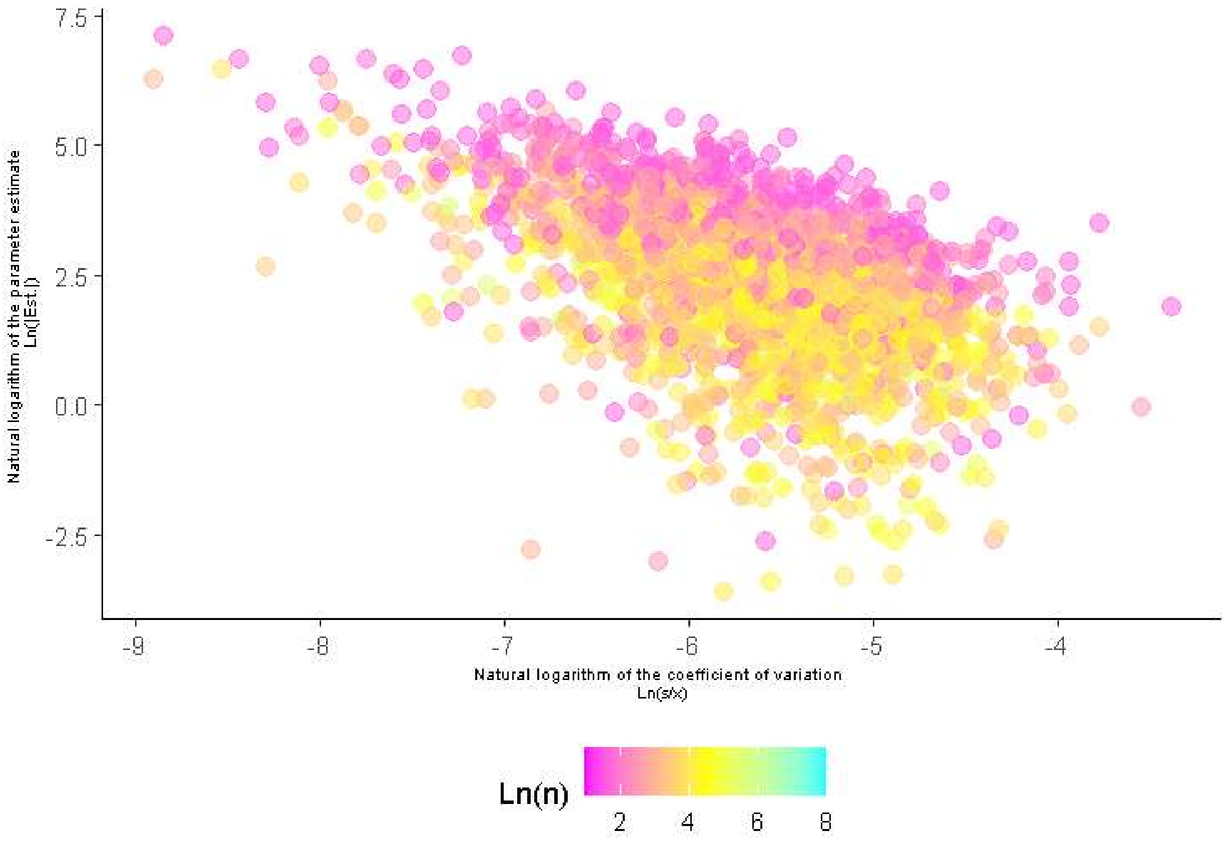


*Figure S3: Simulated values according to the formal principle laid out above.*
